# Impact of *Starmerella bacillaris* and *Zygosaccharomyces bailii* on ethanol reduction and *Saccharomyces cerevisiae* metabolism during mixed wine fermentations

**DOI:** 10.1101/2021.11.11.467398

**Authors:** Angela Capece, Angela Pietrafesa, Gabriella Siesto, Rocchina Pietrafesa, Víctor Garrigós, Patrizia Romano, Emilia Matallana, Agustín Aranda

## Abstract

The bulk of grape juice fermentation is carried out by the yeast *Saccharomyces cerevisiae*, but non-*Saccharomyces* yeasts can modulate many sensorial aspects of the final products in ways not well understood. In this study, some of such non-conventional yeasts were screened as mixed starter cultures in a fermentation defined medium in both simultaneous and sequential inoculations. One strain of *Starmerella bacillaris* and another of *Zygosaccharomyces bailii* were chosen by their distinct phenotypic footprint and their ability to reduce ethanol levels at the end of fermentation, particularly during simultaneous vinification. *S. bacillaris* losses viability strongly at the end of mixed fermentation, while *Z. bailii* remains viable until the end of vinification. Interestingly, for most non-*Saccharomyces* yeasts, simultaneous inoculation helps for survival at the end of fermentation compared to sequential inoculation. *S. cerevisiae* viability was unchanged by the presence of the either yeast. Characterization of both strains indicates that *S. bacillaris* behavior is overall more different from *S. cerevisiae* than *Z. bailii. S. bacillaris* has a less strict glucose repression mechanism and molecular markers like catabolite repression kinase Snf1 is quite different in size. Besides, *S. cerevisiae* transcriptome changes to a bigger degree in the presence of *S. bacillaris* than when inoculated with *Z. bailii. S. bacillaris* induces the translation machinery and repress vesicular transport. Both non-*Saccharomyces* yeast induce *S. cerevisiae* glycolytic genes, and that may be related to ethanol lowering, but there are specific aspects of carbon-related mechanisms between strains: *Z. bailii* presence increases the stress-related polysaccharides trehalose and glycogen while *S. bacillaris* induces gluconeogenesis genes.

## 1. Introduction

Producing wine from grape juice in a traditional way relies on spontaneous fermentation carried out by the microorganisms present on the grape surface and the winery equipment (Ribéreau-Gayon et al., 2006). That implies a complex ecological environment where first yeast, filamentous fungi and bacteria coexist, next only some species of yeasts adapted to the acidic and low oxygen environment proliferate and at the end species of *Saccharomyces* take over due to their high fermentative power and tolerance to ethanol to complete fermentation. The non-*Saccharomyces* (NS), so-called non-conventional yeasts, despite their low fermentative power, contribute greatly to the final product (Jolly et al., 2014)(Benito et al., 2019), as these yeasts produce specific metabolites affecting the wine aroma, or produce enzymes that help liberating volatile compound from the grape (Capece and Romano, 2019). The use of non-*Saccharomyces* yeasts as starters might satisfy an additional request of winemakers, as it could be a potential tool for the reduction of alcohol content in wine. Nowadays, the increase of alcohol levels in wine is one of the main challenges affecting the winemaking sector, due to global climate change which determined an increase of grape maturity (De Orduna, 2010). In this context, the interest for reduction of ethanol content in wine was increased and among the available tools addressed to this aim, the microbiological approach appears a promising way. In particular, researchers’ interest was addressed to investigate the wide variability in ethanol yield among non-*Saccharomyces* yeasts, that could be a potential tool for the reduction of alcohol content in wine (Gonzalez et al., 2013) (Contreras et al., 2015). Low ethanol yield for instance was found in mixed fermentations with some strains belonging to *Hanseniaspora*, and *Zygosaccharomyces* (Gobbi et al., 2014) and *Starmerella* (Milanovic et al., 2012) genera.

Modern enology has developed pure yeast starters to obtain a more reliable fermentation. The usual way to delivered them is in the form of active dry yeast (Pérez-Torrado et al., 2015). Inoculation with strains of *S. cerevisiae* make the whole fermentation process more robust and reliable, but it lacks some of the richness that more microbiologically complex fermentations have. This aspect contributed to an increased interest on the use of non-*Saccharomyces* yeasts in winemaking (Benito et al., 2019)(Padilla et al., 2016) and in the latest years NS yeasts have been added to the portfolio of yeast manufacturers, so dry starters for *Torulaspora, Metschnikowia, Kluyveromyces, Wicheranomyces, Lanchacea* and *Schizosaccharomyces* are marketed [7]. Furthermore, some strains that are not so widespread in the market have interesting winemaking properties, but they need more physiological characterization.

Apiculate yeasts *Hanseniaspora* are the main species present on mature grapes and they produce enzymes and aroma compounds that expand the diversity of wine color and flavor (Martin et al., 2018). A strain of *Hanseniaspora uvarum* is able to reduce ethanol in mixed fermentations (Gobbi et al., 2014). *Starmerella bacillaris* (former *Candida zemplinina*) is an interesting species for the enological point of view. It is a fructophilic yeast and a high glycerol producer, but it produces low ethanol after fermentation, both alone and in mixed fermentation (Englezos et al., 2016)(Lemos Junior et al., 2021). It produces high alcohols, such as benzyl alcohol that inhibit fungal growth. However, it is only produced as cream yeast (Roudil et al., 2019). *Zygosaccharomyces* have been regarded as spoilage yeasts due to their high tolerance to osmotic and acidic stress (Escott et al., 2018). *Zygosaccharomyces* found in grapes and musts are able to increase the production of higher alcohol and reduce acetoin. *Z. bailii* produced wine with reduced ethanol concentration (Contreras et al., 2015)(Zhu et al., 2020).

Since most non-*Saccharomyces* yeasts are unable of completing alcoholic fermentation, *S. cerevisiae* strains should be added in simultaneous or sequentially inoculum modality. When planning mixed fermentations, simultaneous inoculation is the simplest way to proceed, but sequential inoculation (first NS and then *Saccharomyces*) give time to the non-conventional yeast to contribute to the final product with no competitions from *Saccharomyces*. Grape must fermentation, even with just one yeast strains inoculated, is a complex ecological system. There are many kind of interactions, both biotic and abiotic, that link the performance of wine yeasts (Ciani et al., 2016). The most obvious of these interaction is the competition for the nutrients of the grape juice, usually the less abundant ones, like amino acids and vitamins (Medina et al., 2012). More direct interactions rely in the production of toxic molecules, like killer factors, antimicrobial peptides and medium-chain fatty acids. Furthermore, direct cell-to-cell interactions have been linked to the early death of *T. delbrueckii, K. thermotolerans* and S. *bacillaris* by *S. cerevisiae* (Renault et al., 2013)(Nissen et al., 2003)(Englezos et al., 2019b). All those interactions are crossed, so we need more information on the molecular causes behind the metabolic relationships between yeasts that coexist during fermentation.

In this work, a series of potential interesting wine species, were screened in a defined medium for the use as mixed starters with *S. cerevisiae*, by exploring different inoculation regimes in order to establish the inoculum conditions and mixed starter culture that enable the production of wine with reduced ethanol concentration. Two good ethanol-reducing strains (a *S. bacillaris* and a *Z. bailii*) that have an overall impact in metabolites production during fermentations were chosen for a deep analysis of strain physiology. *S. bacillaris* dies at a highest rate than *Z. bailii* in such fermentations. The physiology of those yeasts in selective media happened to be quite different, so the impact of them in the *S. cerevisiae* transcriptomic during mixed fermentations in synthetic grape juice was evaluated to see common and differential impacts. *S. bacillaris* behavior is less similar to *S. cerevisiae* than *Z. bailii*, and cause a greater change in *S. cerevisiae* transcription. Both yeast induce glycolysis, but S. *bacillaris* up-regulates also gluconeogenesis and *Z. bailii* the synthesis of stress polysaccharides like trehalose and glycogen.

## 2. Materials and methods

### 2.1. Yeast strains

Eight non-*Saccharomyces* yeast strains, belonging to UNIBAS Yeast Collection (UBYC), University of Basilicata (Potenza, Italy), were tested in the first step of this study. The tested strains were the following: one *Debaryomyces polymorphus* (Db2), two *Hanseniaspora uvarum* strains (H3 and H9), four *Starmerella bacillaris* strains (St1, St2, St5 and St8) and one *Zygosaccharomyces bailii* (Zb1). All the strains were previously isolated during spontaneous lab-scale fermentations of grape of different varieties, directly collected in the vineyard, or fruit (prickly pear). These strains were identified by restriction analysis of the amplified ITS region (Granchi et al., 1999); the results of restriction analysis were confirmed by analysis of ITS sequences, which were then blasted with NCBI database. In addition, one commercial *S. cerevisiae* strain, EC1118 (Lallemand Inc.), was used.

### 2.2. Stress and growth tests

Strain tolerance to ethanol and SO_2_, expressed as the ratio between the growth in microplates in broth with (14% v/v ethanol and 150 mg/L SO_2_ respectively) and without the stress factor. β-glucosidase and β-xylosidase were measured as previously described (Manzanares et al., 2000)(Manzanares et al., 1999). For the spot analysis, stationary cultures grown in YPD medium (2% glucose, 2% peptone, 1% yeast extract) overnight were serially diluted and 5 μl drops were spotted in YPD, YPS (changing glucose by 2% sucrose), YPG (2% glycerol), SD (25 glucose, 0.17% Yeast Nitrogen Base, 0.5% ammonium sulfate) or SPro (SD with 0.5% proline as nitrogen source). Glucose analog 2-deoxyglucose (2DG) was added to the YPS plates at 200 ng/ml for glucose repression tests. For the H_2_O_2_ oxidative stress test, the equivalent of 2 OD_600_ of overnight YPD cultures were spread on YPD plates and paper circles containing 5 circles containing 5 μl of 30% H_2_O_2_ was placed in the middle and diameter of inhibition halos were measured the following day. To test growth in molasses, the biomass propagation experiments were performed in molasses medium (Torrellas et al., 2020) diluted to 60 g/L sucrose. Cells were cultivated at 30 C with shaking (180 rpm) and growth was followed by OD_600_ and ethanol determined enzymatically with a kit (Megazyme International Ireland Ltd).

### 2.3. Mixed fermentations

The selected strains were tested in laboratory scale fermentations using synthetic grape juice medium (Fleet, 1993), indicated in the OIV-OENO 370-2012 resolution. The final concentration of sugars was 230 g/L (115 g/L of glucose and 115 g/L of fructose), pH adjusted to 3.5. The fermentations were carried out in sterile 130 mL flasks filled with 100 mL of synthetic must, equipped with stoppers and kept under static conditions at 26°C. The flasks were inoculated with 48-h pre-cultures grown in YPD broth at 26°C with shaking. Each non-*Saccharomyces* strain was inoculated in combination with the *S. cerevisiae* strain in two different modalities, sequential (SeF) and simultaneous (SiF) inoculum. In the SeF trials, the inoculum ratio was 1:1, with an inoculation ratio of 1×10^7^ cells/mL for both the strains, but the non-*Saccharomyces* strain was inoculated at time 0, whereas the *S. cerevisiae* strain was added when the alcohol content reach about 5% (v/v). In the SiF trials, the two strains were simultaneously inoculated, but at different inoculum levels (1×10^3^ cells/mL for *S. cerevisiae* strain and 1×10^7^ cells/mL for the non-*Saccharomyces* strain). *S. cerevisiae* EC1118 at concentration of 1×10^7^ cells/mL was used as control.

The fermentation kinetics were monitored daily by measuring the weight loss of the flasks (due to the carbon dioxide release) and sugar consumption. The kinetic growth of yeast strains was checked by plate counting of fermenting must samples on two different agar media, Wallerstein Laboratory (WL) Nutrient Agar medium (Sigma-Aldrich) (Pallmann et al., 2001) and Lysine Agar medium (Oxoid Unipath Ltd, Hampshire, UK) with addition of bromocresol green. Dilution plates containing a statistically representative number of colonies were counted.

### 2.4. Analytical Determinations

Experimental wines obtained from the inoculated fermentation were analyzed for conventional chemical parameters, such as ethanol, total acidity, malic and lactic acid, volatile acidity, residual sugars, glucose, fructose, pH, by a Fourier Transfer Infrared WineScan instrument (OenoFoss™, Hillerød, Denmark). The content of the main secondary influencing wine aroma, such as acetaldehyde, n-propanol, isobutanol, amyl alcohols, ethyl acetate and acetoin, were determined by direct injection gas chromatography of 1 μl sample into a 180 cm × 2 mm glass column packed with 80/120 Carbopack B/5% Carbowax 20 M (Supelco, Bellefonte, PA). The column was run from 70 to 140 °C, the temperature being ramped up at a rate of 7 °C/min. The carrier gas was helium at a flow rate of 20 ml/min. Levels of the secondary compounds were determined by calibration lines, as described by Capece et al. (Capece et al., 2013). Analysis of variance (ANOVA) was used to evaluate differences in chemical compounds of the experimental wines obtained by different inoculation modalities, by using Tukey’s test to compare the mean values. Principal component analysis (PCA) was carried out on the data of wines produced from mixed starters at laboratory scale. The PAST3 software ver. 3.20 (Hammer, Harper, & Ryan, 2018) was used for the statistical analyses.

### 2.5. Western blot

To analyze Snf1 activation and peroxiredoxin status, proteins were extracted by fast cell lysis with trichloroacetic acid (TCA) (Orlova et al., 2008). 5.5% TCA was added to 5 OD_600_ units of cells, the mix was incubated on ice for 15 min and centrifuged. The pellet was washed with acetone twice and resuspended in 150 μL of 10 mM Tris-HCl, pH 7.5, 1 mM EDTA, and broken with 150 μL of 0.2 M NaOH. SDS-PAGE was carried out in an Invitrogen Novex mini-gel device, gel was blotted onto PVDF membranes a Novex semy dry blotter (Invitrogen). The membrane was probed with either anti-AMPKα (Thr172, Cell Signalling Technologies), anti 2-Cys-Prx (Abcam) or anti-Prx-SO_3_ (abcam) antibodies. Anti-actin (Sigma) was used as loading control. The ECL Western blotting detection system (GE) was used following the manufacturer’s instructions in a ImageQuant LAS500 (GE).

### 2.6. Transcriptomic analysis

Fermentation were carried out in synthetic must, inoculating simultaneously *S. cerevisiae* always at 1×10^6^ cells/mL and either non-*Saccharomyces* at 1×10^7^ cells/mL. Cells were taken after 24 h of growth. Total RNA was isolated from yeast pellet using the RNeasy mini kit (Qiagen) following the manufacturer’s instructions for yeast. RNA quality was assessed by using TapeStation System 4200 (Agilent Technologies, Santa Clara, CA, USA). All samples presented an RNA Integrity Number (RIN) value ≥ 8.00. Libraries were prepared using 1 µg of RNA with TruSeq® stranded mRNA Preparation Kit (Illumina, San Diego, CA, USA) followed by sequencing with a Novaseq 6000 sequencer using 150 base read lengths in paired-end mode (Illumina, San Diego, CA, USA) according to manufacturer’s protocol in the facilities of ADM LifeSequencing (Paterna, Spain). Bcl format files were processed to transform them to FASTQ format files using the BCL2FASTQ (version 2.20) software. The reads were trimmed based on their quality, (threshold < Q20) with bbmap software. Adapters and duplicated reads produced by Illumina sequencing were furthermore trimmed with cutadapt v3.0. Clean sequences were mapped against EC1118 Saccharomyces cerevisiae reference genome with SALMON software (Patro et al., 2017). This software maps against the reference genomes and creates a count matrix that will be used in further steps. Differential expression was measured using DESeq2 software (Love et al., 2014). Enrichment functional analysis of the DEG was carried out using gene ontology (GO) databases implemented in the GENEONTOLOGY api (Mi et al., 2019) which performs enrichment by Fisher’s Exact Test and corrects by FDR.

## 3. Results

### 3.1. Characterization of eight non-*Saccharomyces* strains by mixed fermentations: sequential and co-inoculations

Eight wild non-*Saccharomyces* strains, belonging to different species and coded with Db2 (*Debaryomyces polymorphus*), Ha3, Ha9 (*Hanseniaspora uvarum*), St1, St2, St5, St8 (*Starmerella bacillaris*) and Zb1 (*Zygosaccharomyces bailli*), were tested in the first step of the research activity. Table 1 shows the characteristics of enological interest of the selected yeast strains, like enzyme activities and stress tolerance. *H. uvarum* (Ha3, Ha9) and *D. polymorphus* (Db2) have high β-glycosidase and β-xylosidase activities, *Z. bailii* (Zb1) has medium levels, while *S. bacillaris* strains (St1, St2, St5, St8) showed low levels of enzymatic activities, only a low β-glucosidase activity was exhibited by St1 and St2 strains and none β-xylosidase whatsoever. Regarding the other technological parameters, high ethanol tolerance was found in *D. polymorphus* and *Z. bailii*, while *H. uvarum* and *S. bacillaris* strains were more sensitive to this compound. Only the *D. polymorphus* strain showed a low level of sulphur dioxide tolerance, while the rest, as expected for non-*Saccharomyce*s yeasts, are very sensitive to this chemical.

**Table 1.** Main technological characteristics of the selected non-*Saccharomyces* strains.

| Strain code | Quantitative enzymatic activities |  | EtOH<sup>(b)</sup> | SO<sub>2</sub><sup>(b)</sup> |
| --- | --- | --- | --- | --- |
|  | β-glucosidase<sup>(a)</sup> | β-xylosidase<sup>(a)</sup> |  |  |
| Db2 | 157.162 ± 25.98 | 108.182 ± 20.20 | 0.43 ± 0.07 | 0.11 ± 0.01 |
| Ha3 | 116.345 ± 14.43 | 106.141± 5.77 | 0.04 ± 0.00 | 0.01 ± 0.00 |
| Ha9 | 177.570 ± 14.43 | 51.039 ± 2.89 | 0.06 ± 0.01 | 0.01 ± 0.00 |
| St1 | 26.550 ± 8.66 | 0.00 ± 0.00 | 0.09 ± 0.01 | 0.03 ± 0.00 |
| St2 | 8.182 ± 5.77 | 0.00 ± 0.00 | 0.08 ± 0.01 | 0.02 ± 0.00 |
| St5 | 0.00 ± 0.00 | 0.00 ± 0.00 | 0.08 ± 0.02 | 0.03 ± 0.01 |
| St8 | 0.00 ± 0.00 | 0.00 ± 0.00 | 0.08 ± 0.00 | 0.02 ± 0.00 |
| Zb1 | 83.692 ± 8.66 | 22.468 ± 8.66 | 0.62 ± 0.05 | 0.03 ± 0.00 |

<sup>a</sup>Enzymatic activities reported as nmol pNP/mL/h
<sup>b</sup>Strain tolerance to ethanol and SO<sub>2</sub>, expressed as the ratio between the growth in broth with (14% v/v ethanol and 150 mg/L SO<sub>2</sub>) and without the stress factor.

Strains were then tested as mixed starters with a commercial *S. cerevisiae* strain in a defined, standardized synthetic grape juice that will make easier transcriptomic analysis (see below). In this step, the fermentative behavior of mixed starter cultures was tested by using two modality of inoculation, simultaneous and sequential inoculum. The fermentation kinetics, represented by CO_2_ release, of mixed cultures in micro-vinification trials are showed in Fig 1. Regarding simultaneous fermentation (Fig. 1A), *S. cerevisiae* alone is the fastest one, as expected. Next, all three *S. bacillaris* strains behave in a similar way, starting fermentation at day 1. The rest of strains have a slower pace, being *H. uvarum* and *D. polymorphus* quite similar. All the simultaneous fermentations were completed in 14 days, without significant differences among them, except for the mixed fermentation with Zb1+Sc starter, which was completed in 16 days. At the end of the process, the maximum CO_2_ production was found in the fermentation inoculated with St1+Sc starter (about 13.01 g/100 mL), whereas the lowest amount (about 11.99 g/100 mL) was detected in the fermentation inoculated with Ha9+Sc starter. A similar trend for CO_2_ production was observed in sequential fermentations (Fig. 1B), with constant increase in CO_2_ production from the first fermentation day for all non-*Saccharomyces* strains, although clearly slower than *S. cerevisiae* alone. After the fourth fermentation day, it was observed a high increase of CO_2_ production, as a consequence of the addition of *S. cerevisiae*. The amount of CO_2_ produced at the end of fermentation was similar in all the trials, ranging between 13.91 and 15.17 g/100 ml, with highest production found for fermentation inoculated with St8+Sc starter. The duration of fermentation was higher for mixed cultures than for single starter culture; in fact, all the mixed starters completed the process in 15 days, with similar trend all the trials, whereas fermentation inoculated with pure culture of *S. cerevisiae* (control fermentation) ends the process at day 11. The evolution of sugar consumption during the mixed fermentations (Supplementary Fig. S1), reflects the same trend observed for CO_2_ production, as expected.

**Fig. 1.**
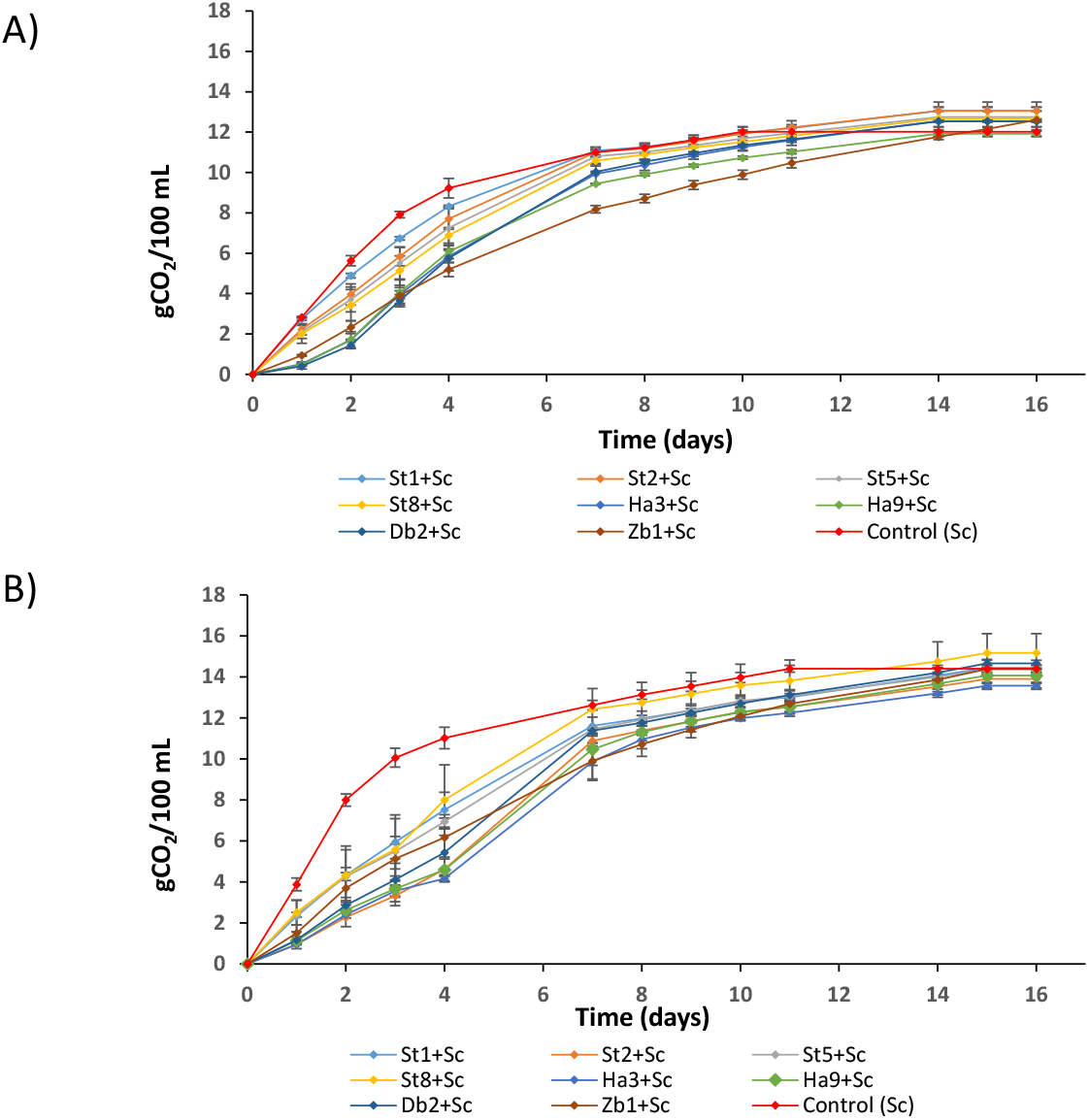
Fermentation kinetics of mixed starters cultures of D. *polymorphus* (Db2), *H. uvarum* (Ha3 and Ha9), *S. bacillaris* (St1, St8, St2 and St5) and *Z. bailii (*Zb1) strains simultaneously A) or sequentially B) inoculated with *S. cerevisiae* EC1118 (Sc). Pure culture of S. cerevisiae EC1118 (Control Sc) was used as a control. Data are means ± standard deviation of two independent experiments.

The studies on yeast population dynamics during inoculated fermentation with mixed starter cultures will help in understanding the interactions between yeast strains (Fig. 2). The persistence level of non-*Saccharomyces* strains during the mixed fermentations was variable in function of yeast strain/species and inoculation modality. In fact, in all the fermentations the presence of non-*Saccharomyces* strains at the end of the process was higher in simultaneous inoculum than sequential inoculum, except for fermentations inoculated with *H. uvarum* strains, in which no differences between two modalities of inoculum were found. In mixed fermentations inoculated with *D. polymorphus* strain (Fig. 2A), in simultaneous modality cells decrease steadily, while in the sequential inoculum, Db2 strain reached a maximum of yeast cells (1.6 × 10^8^ UFC/mL) in 3 days, after that the viable count decreased and at the 10th day of the fermentation no *D. polymorphus* cells were found. The evolution of yeast cells of *H. uvarum* strains (Ha3 in Fig. 2B, Ha9 in Supplementary Figure 2) during the process followed the same trend in both the inoculum modalities, with a decrease of yeast cells after 4 days of fermentation. Furthermore, after 10 days of fermentation, no *H. uvarum* colonies were found on plates. In mixed fermentations with *S. bacillaris* strains (St8 in Fig. 2C, rest in Supplementary Fig. 2), for simultaneous inoculum a slight increase of cell count was observed in the first two days of fermentations, after that cell count slightly was reduced and at the end of the process a number of viable cells ranging between 2 × 10^2^ and 3.4 × 10^3^ UFC/mL was found. Regarding fermentations inoculated in sequential modality, for St8 (Fig. 2C) and St5 (Supplementary Figure 2) strains, in the first days of fermentation a trend similar to simultaneous inoculum was found, with an increase of viable cells, whereas after the third fermentation day the number of viable cells decreased and at the end of the process no *S. bacillaris* cells were found. Regarding mixed fermentation with *Z. bailii* (Figure 2D), in simultaneous inoculum, the number of Zb1 cells remains constant in the first two days of the process, after that a reduction in number of viable cells was observed, although a number quite high of viable cells was found at the end of the fermentation (6 × 10^6^ cells/mL). For sequential inoculum, during the first four days, *Z. bailii* cells increased, after that a high reduction of number of viable cells is observed and in the final wine the number of *Z. bailii* cells was about 1 × 10^5^ cells/mL. As regards the evolution of *S. cerevisiae* EC1118 population in mixed fermentations, similar cell count was observed in both the inoculum modalities and the evolution of *S. cerevisiae* cells in pure culture, used as control, reflects the typical growth kinetic, with the presence of high cell number until the end of the fermentations.

**Fig. 2.**
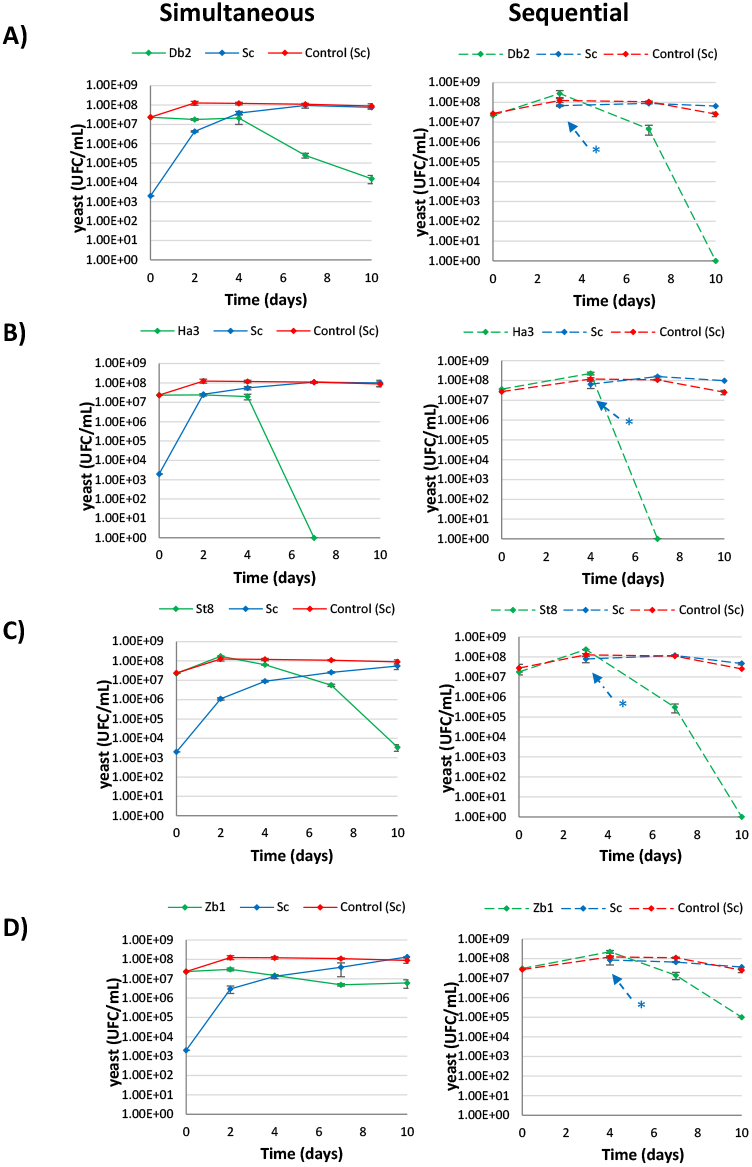
Evolution of yeast populations in mixed fermentations inoculated with *S. cerevisiae* (Sc) and *D. polymorphus* (Db2, A)), *H. uvarum* (Ha3, B)), *S. bacillaris* (St8, C)) and *Z. bailii* (Zb1, D)) in simultaneous and sequential modalities. Values are mean of two independent duplicates. Pure culture of *S. cerevisiae* EC1118 (Control Sc) was used as a control. * indicate the inoculation time of *S. cerevisiae*.

### 3.2. Analysis of experimental wines obtained from mixed starter cultures

The experimental wines were analysed for oenological parameters and main volatile compounds and the data are shown in Tables 2 (simultaneous trials) and 3 (sequential trials). As already reported, all the starter cultures completed the fermentations. Regarding co-inoculum, all the samples from mixed fermentations contained an ethanol concentration lower than control sample (Figure 3A), with values ranging from 11.58 to 12.19 % (v/v), whereas the ethanol content of wine from single fermentation was 12.38 % (v/v). As regards volatile acidity, the activity of non-*Saccharomyces* strains did not increase it; in fact, the volatile acidity of samples inoculated with mixed starters was lower or very similar to level detected in control wine, except for the mixed fermentations inoculated with Db2+Sc and Ha3+Sc starters. The experimental wines were analysed also for the content of volatile compounds usually present in high quantity in wines, and involved in the wine flavour. The content of acetaldehyde ranged between 38.72 mg/L (Zb1+Sc) and 65.95 mg/L (Ha9+Sc), being the latter the only case where more acetaldehyde is produced compared with the control. The ethyl acetate was found in concentrations ranging from 7.71 mg/L (St8+Sc) to 28.95 mg/L (Db2+Sc), with higher levels than the control when *D. polymorphus* and *H. uvarum* were used. Generally, mixed starters produced experimental wines characterized by lower amount of alcohols (n-propanol, isobutanol, amyl alcohols) than the wine from *S. cerevisiae* EC1118 strain, used as control, except for n-propanol. The highest difference between single and mixed starter wines was found for fermentation performed by mixed starter including *S. bacillaris* strains, which contained lower amounts of both D-amyl and isoamyl alcohols than experimental wine fermented with EC1118 strain.

**Table 2.**
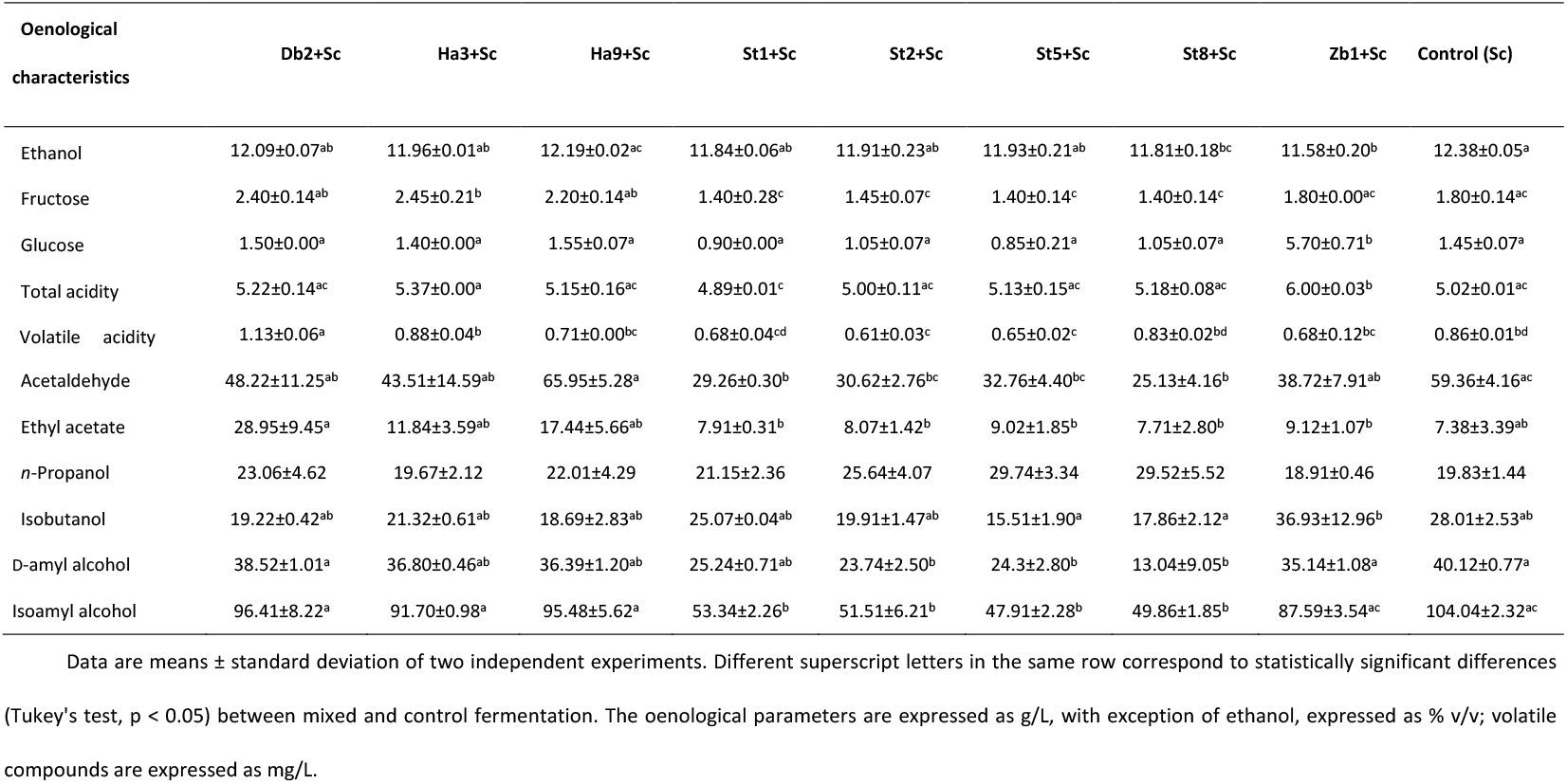
Oenological parameters and main volatile compounds of experimental wines obtained from mixed starters cultures of selected non-*Saccharomyces* strains (Db2, Ha3, Ha9, St1, St8, St2, St5 and Zb1) simultaneously inoculated with *S. cerevisiae* EC1118 (Sc). Pure culture of *S. cerevisiae* EC1118 (Control Sc) was used as control.

| Oenological characteristics | Db2+Sc | Ha3+Sc | Ha9+Sc | St1+Sc | St2+Sc | St5+Sc | St8+Sc | Zb1+Sc | Control (Sc) |
| --- | --- | --- | --- | --- | --- | --- | --- | --- | --- |
| Ethanol | 12.09±0.07<sup>ab</sup> | 11.96±0.01<sup>ab</sup> | 12.19±0.02<sup>ac</sup> | 11.84±0.06<sup>ab</sup> | 11.91±0.23<sup>ab</sup> | 11.93±0.21<sup>ab</sup> | 11.81±0.18<sup>bc</sup> | 11.58±0.20<sup>b</sup> | 12.38±0.05<sup>a</sup> |
| Fructose | 2.40±0.14<sup>ab</sup> | 2.45±0.21<sup>b</sup> | 2.20±0.14<sup>ab</sup> | 1.40±0.28<sup>c</sup> | 1.45±0.07<sup>c</sup> | 1.40±0.14<sup>c</sup> | 1.40±0.14<sup>c</sup> | 1.80±0.00<sup>ac</sup> | 1.80±0.14<sup>ac</sup> |
| Glucose | 1.50±0.00<sup>a</sup> | 1.40±0.00<sup>a</sup> | 1.55±0.07<sup>a</sup> | 0.90±0.00<sup>a</sup> | 1.05±0.07<sup>a</sup> | 0.85±0.21<sup>a</sup> | 1.05±0.07<sup>a</sup> | 5.70±0.71<sup>b</sup> | 1.45±0.07<sup>a</sup> |
| Total acidity | 5.22±0.14<sup>ac</sup> | 5.37±0.00<sup>a</sup> | 5.15±0.16<sup>ac</sup> | 4.89±0.01<sup>c</sup> | 5.00±0.11<sup>ac</sup> | 5.13±0.15<sup>ac</sup> | 5.18±0.08<sup>ac</sup> | 6.00±0.03<sup>b</sup> | 5.02±0.01<sup>ac</sup> |
| Volatile acidity | 1.13±0.06<sup>a</sup> | 0.88±0.04<sup>b</sup> | 0.71±0.00<sup>bc</sup> | 0.68±0.04<sup>cd</sup> | 0.61±0.03<sup>c</sup> | 0.65±0.02<sup>c</sup> | 0.83±0.02<sup>bd</sup> | 0.68±0.12<sup>bc</sup> | 0.86±0.01<sup>bd</sup> |
| Acetaldehyde | 48.22±11.25<sup>ab</sup> | 43.51±14.59<sup>ab</sup> | 65.95±5.28<sup>a</sup> | 29.26±0.30<sup>b</sup> | 30.62±2.76<sup>bc</sup> | 32.76±4.40<sup>bc</sup> | 25.13±4.16<sup>b</sup> | 38.72±7.91<sup>ab</sup> | 59.36±4.16<sup>ac</sup> |
| Ethyl acetate | 28.95±9.45<sup>a</sup> | 11.84±3.59<sup>ab</sup> | 17.44±5.66<sup>ab</sup> | 7.91±0.31<sup>b</sup> | 8.07±1.42<sup>b</sup> | 9.02±1.85<sup>b</sup> | 7.71±2.80<sup>b</sup> | 9.12±1.07<sup>b</sup> | 7.38±3.39<sup>ab</sup> |
| *n*-Propanol | 23.06±4.62 | 19.67±2.12 | 22.01±4.29 | 21.15±2.36 | 25.64±4.07 | 29.74±3.34 | 29.52±5.52 | 18.91±0.46 | 19.83±1.44 |
| Isobutanol | 19.22±0.42<sup>ab</sup> | 21.32±0.61<sup>ab</sup> | 18.69±2.83<sup>ab</sup> | 25.07±0.04<sup>ab</sup> | 19.91±1.47<sup>ab</sup> | 15.51±1.90<sup>a</sup> | 17.86±2.12<sup>a</sup> | 36.93±12.96<sup>b</sup> | 28.01±2.53<sup>ab</sup> |
| D-amyl alcohol | 38.52±1.01<sup>a</sup> | 36.80±0.46<sup>ab</sup> | 36.39±1.20<sup>ab</sup> | 25.24±0.71<sup>ab</sup> | 23.74±2.50<sup>b</sup> | 24.3±2.80<sup>b</sup> | 13.04±9.05<sup>b</sup> | 35.14±1.08<sup>a</sup> | 40.12±0.77<sup>a</sup> |
| Isoamyl alcohol | 96.41±8.22<sup>a</sup> | 91.70±0.98<sup>a</sup> | 95.48±5.62<sup>a</sup> | 53.34±2.26<sup>b</sup> | 51.51±6.21<sup>b</sup> | 47.91±2.28<sup>b</sup> | 49.86±1.85<sup>b</sup> | 87.59±3.54<sup>ac</sup> | 104.04±2.32<sup>ac</sup> |

Data are means ± standard deviation of two independent experiments. Different superscript letters in the same row correspond to statistically significant differences (Tukey's test, $p < 0.05$ ) between mixed and control fermentation. The oenological parameters are expressed as g/L, with exception of ethanol, expressed as % v/v; volatile compounds are expressed as mg/L.

**Fig. 3.**
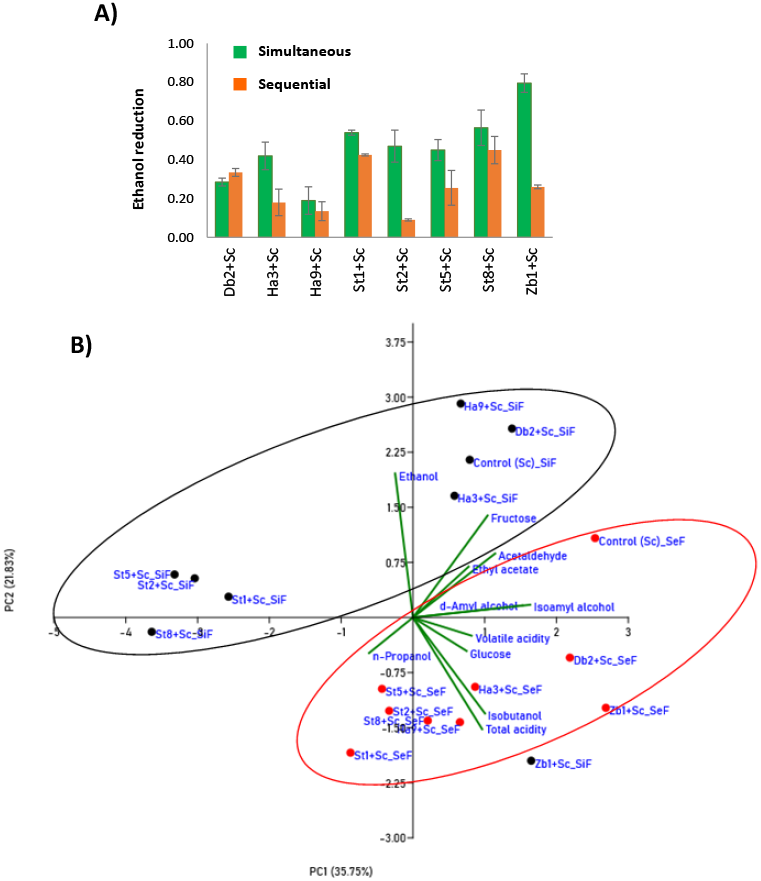
Metabolite analysis of mixed fermentations. A) Starter ability to reduce the ethanol content in experimental wines obtained by simultaneous (green) and sequential (orange) inoculation as ratio between ethanol produced by *S. cerevisiae* pure culture and ethanol produced from each mixed starter. B) Principal component analysis (PCA) biplot of the oenological parameters and main volatile compounds detected in experimental wines obtained by simultaneous (SiF) (black) and sequential (SeF) (red) inoculum of selected non-*Saccharomyces* strains. Pure culture of *S. cerevisiae* EC1118 (Control Sc) was used as control.

The samples obtained by mixed starter sequentially inoculated with EC1118 strain contained lower ethanol than control experimental wine, as already found for co-inoculation (Figure 3A). The lowest level of volatile acidity was found in experimental wine obtained by H9+Sc starter (0.69 mg/L), a result very surprising as apiculate yeasts are known to be high producers of acetic acid, whereas in the other samples volatile acidity ranged between 0.91 and 1.18 mg/L. Regarding acetaldehyde, this compound was detected in the range between 64.38 (Zb1+Sc, well above the control fermentation) and 27.75 mg/L (St1+Sc), whereas the ethyl acetate content ranged from 6.85 mg/L (St1+Sc, below control) to 28.14 mg/L (Db2+Sc, above control). All mixed cultures produced wines containing slightly higher amount of n-propanol, than the wine from *S. cerevisiae* strain. Regarding the levels of the other higher alcohols detected (isobutanol and amyl alcohols), control wine contained higher level of these compounds than samples from mixed starters. Although these compounds represented the most abundant groups in all the analysed samples, all the starters produced an amount of high alcohols lower than 300 mg/L, which is within the acceptable level for these compounds.

The ability of mixed starter to reduce the ethanol content was calculated as ratio between ethanol produced by *S. cerevisiae* pure culture and ethanol produced from each mixed starter (Fig. 3A). All the mixed starters, in both inoculum modalities, determined an ethanol reduction, as already reported. The reduction level was higher for simultaneous than for sequential inoculum for all the starters, except Db2+Sc, in which a slightly higher reduction was found in sequential inoculum. In fermentations using simultaneous inoculations, the ethanol reduction ranged between 0.19 (Ha9+Sc) until 0.80 (Zb1+Sc), whereas in sequential inoculation the starter ability to reduce the ethanol content ranged between 0.09 (St2+Sc) and 0.45 (St8+Sc).

All the parameters determined in the experimental wines were submitted to Principal Component Analysis (PCA). The plot of all the experimental wines on the plane defined by the first two components is shown in Fig. 3B. The two principal components, PC1 and PC2, accounted for 58% of the total variance (36 and 22%, respectively). The PC1 was positively correlated mainly with D-amyl and isoamyl alcohols and negatively mainly associated with n-propanol, whereas the PC2 was mainly positively related to content of ethanol and residual fructose and negatively with total acidity and isobutanol. This analysis allowed to differentiate the experimental wines in function of inoculation modality; in fact, almost all the samples obtained with non-*Saccharomyces* strains inoculated simultaneously with *S. cerevisiae* strain are located in upper part of the scatterplot (except St8+Sc and Zb1+Sc, Figure 3B), whereas all the experimental wines obtained by sequential inoculum are grouped together in the lower part of the scatterplot. The only exception is represented by the two wines obtained with mixed starter including *Z. bailii* strain, which were located very near, in both the inoculation modalities. As expected, the experimental wines obtained with pure culture of EC1118 strain are located in the same quadrant of the plot. Furthermore, as regards the wines obtained by using the different *S. bacillaris* strains, in both the inoculation modalities the experimental wine grouped very near and far from the others, with exception of wine fermented with starter including St8 strain, which was located in a quadrant different from the wines fermented with mixed starters including St1, St2 and St5.

### 3.3. Physiological study of strains of interest

Next, the experiments were focused on two strains, *S. bacillaris* St8 and *Z. bailii* Zb1. These strains were selected as they determined relatively high ethanol reduction during laboratory scale fermentations and a big impact in overall metabolites after fermentation (Fig. 3). To further characterize them, they were tested in different growth conditions (Fig. 4), like different carbon sources (Fig. 4A). In standard, rich, glucose-containing medium YPD, all three strains grow fine, being *Z. baill*i a little bit slower. Similar results were obtained when sucrose (YPS) was used as carbon source. In such medium, glucose repression mechanisms can be tested, using a non-metabolizable glucose analog, 2-deoxyglucose (2DG); in this case a strong repression of growth was observed for *Z. bailli* and also (as expected) in *S. cerevisiae*. Interestingly enough, *S. bacillaris* is immune to this inhibitor, indicating that the mechanism of standard catabolite repression does not work for this yeast. When the carbon source was glycerol, a fully respiratory source, *S. cerevisiae* is the only one that grows well, and the other two strains struggle. This poor respiration may indicate poor management of oxygen radicals and oxidative stress sensitivity, but no high differences in tolerance to the oxidant H_2_O_2_ were found (Fig. 4B). In fact, *S. bacillaris* is slightly better equipped to deal with this stress than *S. cerevisiae*. Last, nitrogen sources were tested. In the standard minimal, ammonium-containing medium SD, all three strains were able to growth. Ammonium is a rich nitrogen source, but neither in poor proline (SPro), there are no big differences, although *S. bacillaris* is slightly delayed compared to ammonia.

**Fig. 4.**
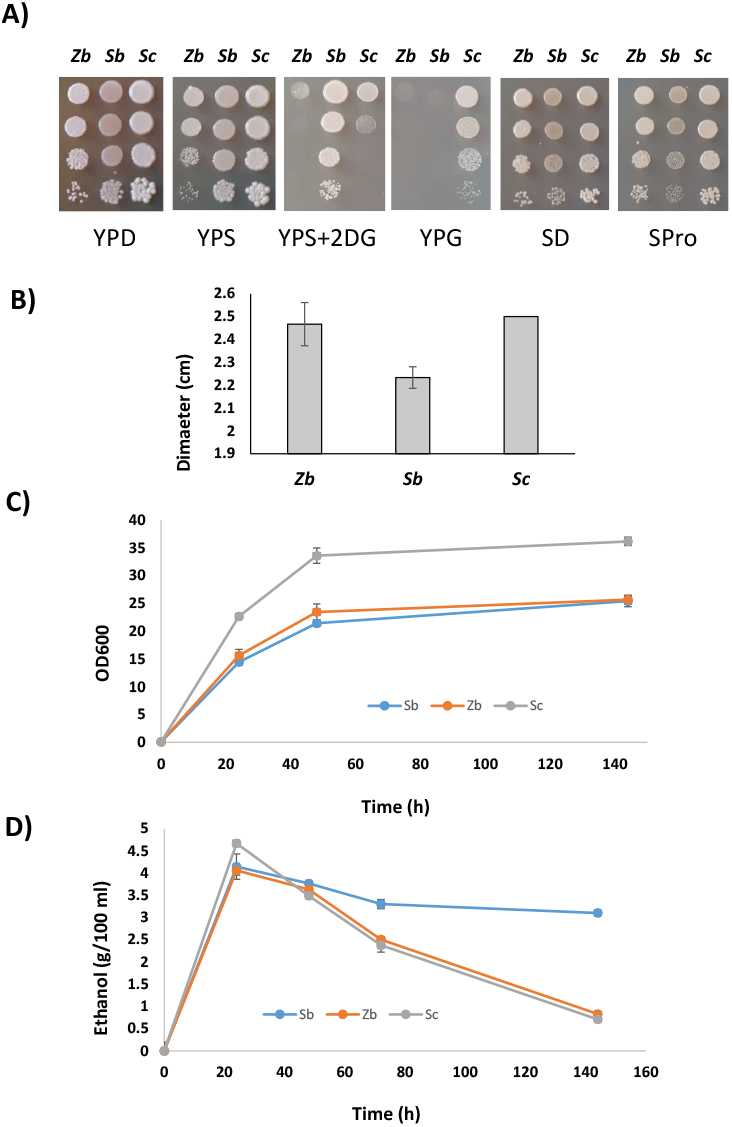
Physiological study of strains of interest. A) Spot analysis of growth on different carbon and nitrogen sources. *Z. bailii* (Zb), *S. bacillaris* (Sb) and *S. cerevisiae* (Sc) strains were grown overnight in YPD liquid to stationary phase, serial dilutions were made and 5 μl drops were spotted in YPD, YPS (sucrose), YPS containing 200 ng/ml 2-deoxyglucose (2DG), YPG (glycerol), SD (minimal medium with ammonium as sole nitrogen source), SPro (minimal medium with proline as sole nitrogen source). B) Oxidative analysis of those strains. Overnight cultures form YPD were plated on YPD plates and a paper circle containing 5 μl drops of hydrogen peroxide was placed in the middle. Inhibition halos were measured the next day. Experiments were carried in triplicate and mean and standard deviation is shown. C) Growth in beet molasses followed by OD_600_ measurement. D) Ethanol measurement from the growth curve showed in panel C). Experiments carried out in triplicates and average and standard deviation is shown.

Another tested condition was the growth of the three strains in sugar beet molasses (Fig. 4C), the media that is used for biomass propagation. In such conditions, *S. cerevisiae* proved to be the fastest one in terms of growth. Both non-*Saccharomyces* yeasts grow slower and reached a lower OD_600_ at saturation point. However, they assimilate sucrose (that is the main sugar in molasses) well, as it disappeared for all strains after 24 h (data not shown). Ethanol produced can give us some ideas about their respiratory metabolism (Fig. 4D). The peak of ethanol was observed early, at the first day, when *S. cerevisiae* produce slightly more ethanol than the other two strains, after that *S. cerevisiae* and *Z. bailli* consume efficiently ethanol during the postdiauxic phase, while *S. bacillaris* does it more inefficiently, indicating again that is the one with a more distant phenotype in metabolic terms.

### 3.4. Molecular analysis of mixed fermentations in synthetic grape juice

Our next goal is to understand the molecular behaviour behind mixed fermentation, analysing gene expression by transcriptomic analysis and performing biochemical analysis of molecular markers of stress response and nutrient signalling. The goal was to evaluate the impact of the non-conventional yeasts on *S. cerevisiae*, so simultaneous inoculation was chosen to allow this interaction from the beginning, plus it induces a bigger ethanol difference (Fig. 3A), but with different inoculation ratios to have more *S. cerevisiae* biomass for RNA and protein extraction. *S. cerevisiae* was inoculated at concentration of 10^6^ cells/ml (higher than before to be able to have enough cells at shorter times), while *S. bacillaris* and *Z. bailli* were inoculated tenfold to be in excess, at 10^7^ cells/ml, so there is an initial excess of non-conventional yeasts, but the amount of *S. cerevisiae* is significant. The overall course of fermentation was followed by CO_2_ production (Fig. 5A) and reducing sugar measurement (Fig. 5B). All fermentations finished, but the mixed fermentations are complete slightly faster than the one with only *S. cerevisiae*, probably due to the smaller inoculum of the latter. Again, experimental wines from mixed fermentation showed lower ethanol as seen in natural grape juice (Fig. 5C). Cells were diluted and spread in selective media to quantify the evolution of each one (Fig. 5D). *S. bacillaris* grows during the first 24 hours, and at this time *S. cerevisiae* reaches a similar cellular density. From that point, the viable numbers of *S. bacillaris* decline, and at 144 h there are no detection of this strain, while *S. cerevisiae* keeps it viability. *Z. bailli* has a different growth profile. It reaches the maximum population level later, at 48 hours, and then the viability is reduced, but it survives longer than *S. bacillaris* did. *S. cerevisiae* with *Z. bailli* grows vigorously and remain high, as was seen with *S. bacillaris*. In fact, both have a similar profile than *S. cerevisiae* alone, so the non-*Saccharomyces* do not offer a serious problem for growth as previously shown. Supplementary Table S1 shows the final concentration of metabolites of such fermentations.

**Fig. 5.**
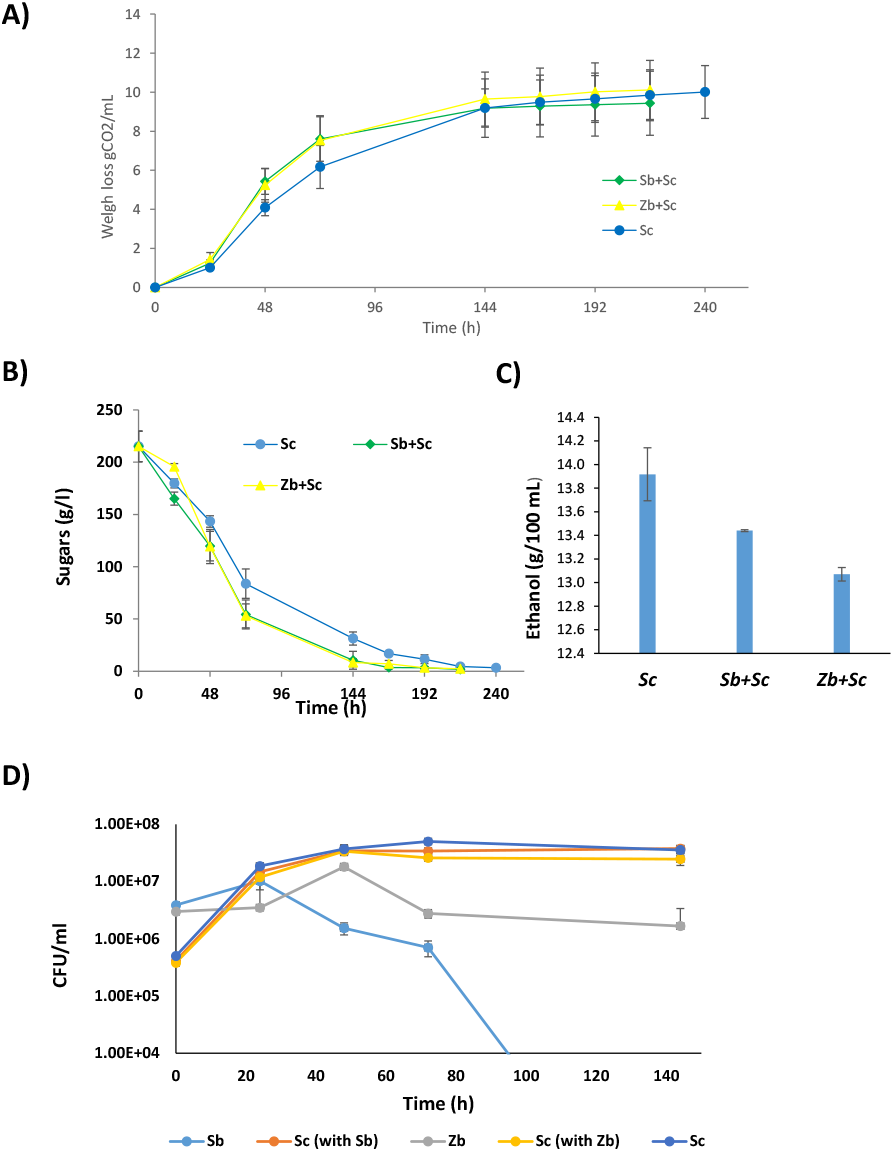
Mixed fermentation of selected strains for molecular analysis. *S. cerevisiae* (Sc) alone and mixed fermentations with *S. bacillaris* (Sb+Sc) or *Z. bailii* (Zb+Sc) were carried out in synthetic grape juice. A) Weight loss of the fermentation. B) Reducing sugars during fermentation. C) Ethanol production at the end of fermentation. D) Cell viability during fermentation.

Samples were taken at day 1, 2 and 3 to extract proteins and to check molecular markers (Fig. 6). First to check for stress response, the oxidative stress protein peroxiredoxin Tsa1 was detected using a specific antibody. A positive control was including consisting in cells grown in rich medium YPD and challenged with oxidative stress in the form of hydrogen peroxide. In those conditions, *S. cerevisiae* and *Z. bailli* showed a clear induction of the protein, indicating that it is reacting to that stress. However, *S. bacillaris* does not produce a reaction to this antibody, so its Tsa1 is not detected in this experiment. In the fermentation with *S. cerevisiae* alone, Tsa1 goes up until 48 h and then its levels are reduced. As *S. bacillaris* does not give signal, the profile in Sc+Sb (*S. cerevisiae* + *S. bacillaris*) comes only from *S. cerevisiae*. In this case, there is a higher level at longer times, indicating a more stressful environment in this mixed fermentation. In Sc+Zb (*S. cerevisiae* + *Z. bailii*) fermentation a strong Tsa1 signal was observed, similar to the oxidative stress cells, so one of them, or both, are in a non-optimal position. There is an antibody that reacts to sulfinilated Tsa1, indicating a hyperoxidaton caused by big stress, but this response was observed in presence of H_2_O_2_, which is not the case of grape juice fermentation, indicating that there is not a strong, irreversible oxidative stress during winemaking, including mixed fermentations. Next, an antibody against the phosphorylated form of AMPK kinase Snf1 was used. That marks the activity of kinase, due to starvation or stress. In this case, *S. bacillaris* gave also signal, but with a different size, in the oxidative stress test. *S. cerevisiae* and *Z. bailli* are again very similar. As seen before (Vallejo et al., 2020), during fermentation, Snf1 is activated very early and its activation decreases later on, at day 3. In the Sc+Sb mixed fermentation the pattern for *S. cerevisiae* is similar than the single fermentation, and in the case of *S. bacillaris’* band, the pattern is similar, but it goes down faster, at day 2. In Sc+Zb, both species cannot be discriminated, but the overall pattern is similar, with early induction and decrease over the next time points. However, an extra band appears suggesting some interactions causing some event of posttranscriptional modification that is altering Snf1 in possible one of those strains. Overall, *S. bacillaris* molecular markers are quite distinctive than the other two species.

**Fig. 6.**
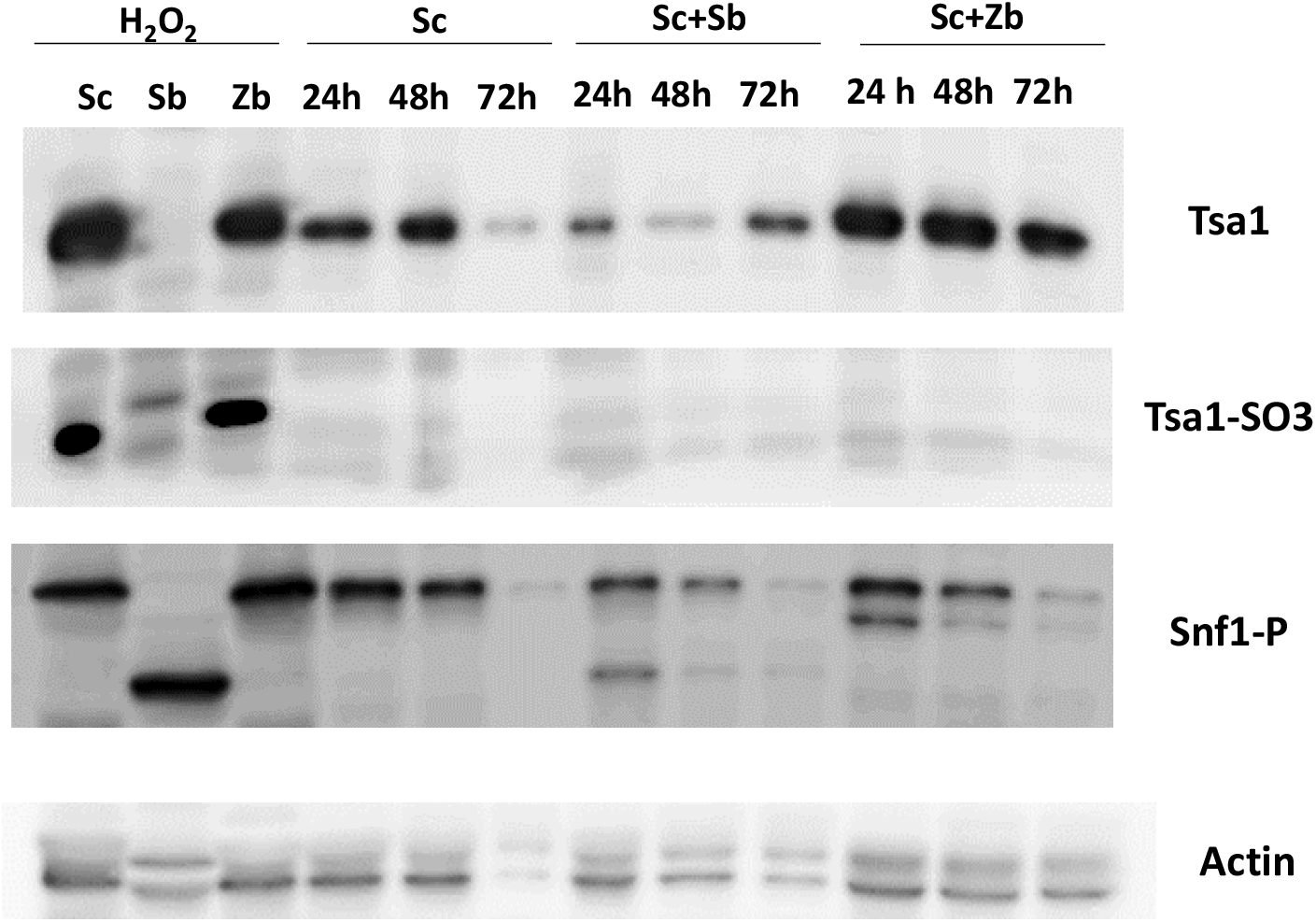
Western blot analysis of peroxirredoxin Tsa1 and glucose repression kinase Snf1 during mixed fermentations. Samples from 1, 2 and 3 days of the fermentation described in Fig. 5 were taken and proteins extracted. A control of oxidative stress was made in cells growing exponentially in YPD challenged with 5 mM H_2_O_2_ for 30 min. Blots with the same amount of proteins were blotted and probed with anti-Tsa1, anti-Tsa1-SO_3_ and Snf1-P specific antibodies. Actin was used as loading control.

### 3.5. Transcriptomic analysis of *S. cerevisiae* under mixed fermentations

Next, a transcriptomic analysis of *S. cerevisiae* gene expression in the presence of the other two yeast species was carried out, in order to see the common and differential impact of *Z. bailii and S. bacillaris*. Considering fermentation advance and growth in synthetic grape juice (Fig. 5), cells were collected at day 1 of fermentation. At this point, *S. cerevisiae* is actively growing and its gene expression machinery should be functioning at maximum capacity, while non-*Saccharomyces* have been present in high numbers and also metabolically active, and the amount of all yeasts is similar (Fig. 5D). Three samples were taken: *Z. bailii*+ *S. cerevisiae, S. bacillaris* + *S. cerevisiae* and *S. cerevisiae* alone. *S. cerevisiae* genes levels were compared among the different samples, giving three comparisons, represented as volcano plots in Fig. 7. At first glance, the presence of *S. bacillaris* (panel 7B) has a bigger impact on *S. cerevisiae* than *Z. bailii* (panel 7A). Fig. 7C shows the genes that are differentially expresses between both mixed fermentations. There are 498 *S. cerevisiae* genes upregulated at least two times in Zb+Sc compared to Sc, and 179 genes downregulated (Supplementary Table S2). Among them, some functional gene ontology categories of biological processes are overrepresented (Table 4 and Supplementary Table S3). Many are involved in metabolism, like “de novo’ NAD biosynthetic process from tryptophan”, “glycolytic process”, “trehalose metabolic process”, “glycogen metabolic process” and many more devoted to energy production and storage. Therefore, it seems that *Z. bailii* is competing with *S. cerevisiae* for resources, causing a general activation of metabolism. Of particular interest is the increase of glycolysis in a fermentative situation. Many glycolytic genes are up-regulated, for instance all for glyceraldehyde-3-phosphate dehydrogenase isozymes, *TDH1, TDH3* and particularly *TDH2*. Alcohol dehydrogenase I *ADH1* is also activated, so that would mean than *S. cerevisiae* is fully producing ethanol, so any alcohol reduction is happening at a different level. “Cortical actin cytoskeleton organization” and “fungal-type cell wall organization” indicate some structural stress in terms of cytoskeleton and cell wall that have to be compensated. There are 179 genes downregulated 2-fold or more. No GO category is over-represented among them, probably because 124 of them are of unknown function, indicating that not so-well defined processes may be in place.

**Fig. 7.**
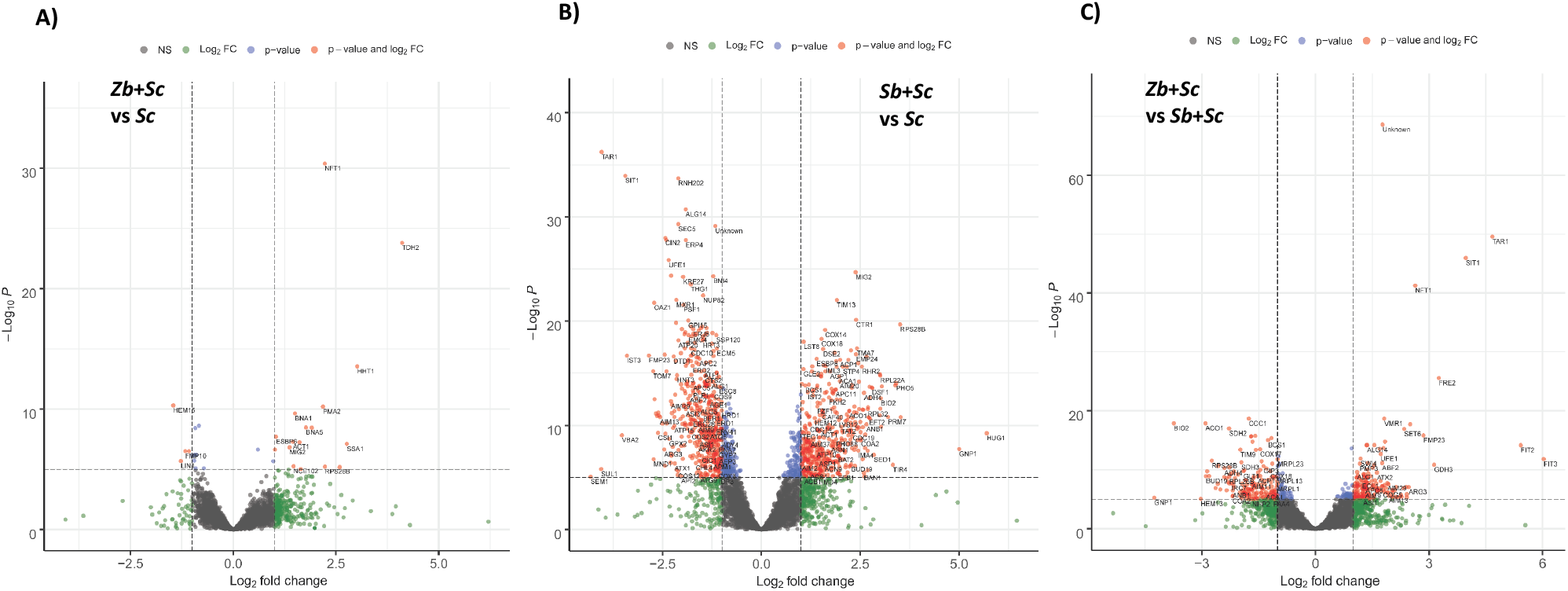
Transcriptomic analysis of mixed fermentations. Volcano plot of: A) *Z*.*bailii* + *S. cerevisiae* fermentation vs *S. cerevisiae* alone control. B) *S. bacillaris*+ *S. cerevisiae* fermentation vs *S. cerevisiae* alone control. C) *Z*.*bailii* + <u>S.cerevisiae</u> fermentation vs *S. bacillaris*+ *S. cerevisiae* fermentation. Log_2_ of fold change and log_2_ of p-value were plotted.

The impact of *S. bacillaris* in numerically *S. cerevisie* is more intense (Fig. 7B). There are 1063 genes relatively up-regulated compared to the control fermentation. Again, genes involved in hexose metabolism, glycolysis/gluconeogenesis, are up regulated, suggesting competition for sugars (Table 2). Many biological function categories are ribosome related, like “rRNA export from nucleus” and “cytoplasmic translation”, indicating that *S. bacillaris* is inducing an increase in protein synthesis in *S. cerevisiae* in a specific way. 1032 genes are downregulated, and interestingly there is overrepresentation of genes involved in intracellular transport, like “cytoplasm to vacuole transport by the Cvt pathway”, “protein targeting to vacuole “, “endoplasmic reticulum to Golgi vesicle-mediated transport” and so on. Therefore, many aspects of the vesicular transport are repressed, so the increase of translation does not seem to be targeted for secretion or membrane targeting.

Next, a direct comparison between *Zb*+*Sc* and *Sb*+*Sc* was made (Fig. 7C). This way, the common effects caused by competition with any yeast will be ruled out, and specific effects will be highlighted. There are 773 *S. cerevisiae* genes that up-regulated in the presence of *Z. bailii* compared to *St. bacillaris* (or down-regulated in *St. bacillaris* compared to *Z. bailii*) (Supplementary Table S2). Conversely, there are 624 genes down-regulated in the presence of *Z. bailii vs St. bacillaris*. Regarding GO analysis, among the categories overrepresented in the up-regulated *S. cerevisiae* genes in the Zb vs Sb “late nucleophagy” and “autophagy of mitochondrion” (Supplementary Table S3) are present, including many *ATG* genes devoted to autophagy. Many of those genes were downregulated in Sb+ Sc vs Sc, so *S. bacillaris* has the ability to repress autophagy processes in *S. cerevisiae*. Amino acid metabolic categories as “homoserine metabolic process”, “cysteine metabolic process” and “methionine biosynthetic process” are also present in the analysis. Finally, many related categories involving ion transport, particularly iron (iron ion transmembrane transport, copper ion transport, cellular iron ion homeostasis…) are present. These involved many siderophores (*ARN1, ARN2, SIT1*), ferric reductases (*FRE1, 2,3* and *5*), iron oxidase (*FET3*) and iron permease (*FTR1*). Siderophore genes are indeed down-regulated in the presence of *S. bacillaris*, while *Z. bailli* has no effect on them. Interestingly, there are two genes of cell wall mannoproteins involved in the retention of siderophore-iron in the cell wall, *FIT2* and *FIT3*, that are repressed in the presence of *S. bacillaris*, but induced in the presence of *Z. bailii*, so they show big differences when data are compared directly (Fig 7C). It seems that *S. bacillaris* provide an excess of iron in the form of siderophores that *S. cerevisiae* sense, reducing their mechanisms of uptake. Last, the 624 genes down-regulated in the presence of *Z. bailii vs S. bacillaris* are enriched in genes involved in rRNA metabolism and translation. The glycolytic genes did not appear in that kind of comparison, so it seems clear that its induction is a common feature in the presence of an alien yeast, probably due to nutrient competition rather than direct interaction.

## 4. Discussion

Complex ecological interactions happen inside spontaneous grape juice fermentations. In this work we tried to analyse such information simplifying the interaction between two yeast species in a standardized growth medium that mimics grape juice, and allow us to perform global analysis, such transcriptomic studies, and protein analysis in a controlled environment. An initial screen using eight different strains and two inoculation strategies (simultaneous and sequential) allowed the selection of strains and conditions. The presence of additional strains other than *S. cerevisiae* always delay fermentation progression, even from the very beginning, when most nutrients are in theory not exhausted and without a decrease in *S. cerevisiae* viability (Fig. 2 and 5). Therefore, metabolism should be affected by the cohabitation with another kind of yeast. Analogous results on the time courses of sequential fermentations were recently found by several authors. Englezos et al. (Englezos et al., 2019a) reported that a sequential fermentation with *S. bacillaris* and *S. cerevisiae* in white grape must took 14 days to finish, while 9 days were needed for the single inoculation with *S. cerevisiae*. Although fast and reliable completion of fermentation are of primary importance in the wine industry, the advantages of the use of non-*Saccharomyces* yeasts in mixed fermentations is thought to compensate the slower fermentation with the advantages in wine quality (Hranilovic et al., 2018). Moreover, a slow fermentation kinetics could be considered as positive for a better retention of volatile compounds (Medina et al., 2013). Our results confirmed that the use of non-*Saccharomyces* strains affected wine characteristics; in fact, the experimental wines from mixed starters differed from single starter wine (Fig. 3). In the case of simultaneous inoculum, the wines obtained by using *S. bacillaris* and *Z. bailii* strains differed more from single starter wine than *H. uvarum* (Fig. 3B). Viability loss could be the cause for *H. uvarum* low impact (Fig. 2). This behaviour could be due to the low competition of these species, which was increased in the case of the delayed inoculation of the *S. cerevisiae* strain, as reported by other authors for different non-*Saccharomyces* species (Wang et al., 2016) (Gobbi et al., 2014). *H. uvarum* is more sensitive to death at the end of fermentation than *S. bacillaris* measured with direct *in situ* fluorescent hybridization (Wang et al., 2014). This phenomenon could have numerous explanations. The loss of viability of the non-*Saccharomyces* in the mixed fermentations can be related to production of yeast metabolites, such as ethanol, medium chain fatty acids and acetaldehyde. Direct cell-to-cell contact has been proved for the pair *S. bacillaris-S. cerevisiae* (Englezos et al., 2019b).

We have focused on two strains, a *S. bacillaris* and a *Z. bailli* for their deeper impact of fermentation and higher ethanol reduction (Fig. 3), in order to get a deeper understanding of the interaction between different yeast species. Both strains were able to reduce the ethanol level of the wine (Tables 2 and 3), but they are very different regarding other parameters. *S. bacillaris* loses viability at the end of fermentation much faster than *Z. bailii* during wine fermentation (Figure 2 and 5), it is insensitive to glucose repression (Fig. 4A) and both are less adapted to respiratory metabolism than *S. cerevisiae* (Fig. 4A). *Z. bailii* viability remains high (Fig. 2), but it is the strain that leads to a slower fermentation (Fig. 1). Therefore, these strains may induce low ethanol production by similar means or by mechanisms completely different. Genetically it is known that *S. bacillaris* lies further away than the two others (Masneuf-Pomarede et al., 2016). Molecular markers of *S. bacillaris* are less similar to *S. cerevisiae* than *Z. bailli* (Fig. 6). Glucose derepression kinase Snf1 is more similar in size between *S. cerevisiae* and *Z. bailli*, while in *S. bacillaris* it is smaller. However, the pattern of induction is quite similar, although targets may differ. Genomic analysis of two strains of *S. bacillaris* suggests a regulation of the fermentation/respiration processes that is partially different from that of *S. cerevisiae* (Lemos Junior et al., 2018), although *SNF1* gene is present. A reason for ethanol reduction could be the respiratory metabolism of non-*Saccharomyces* yeasts. However, *Z. bailli* has been described as a Crabtree positive yeast (Rodicio and Heinisch, 2009) and species of *Z. bailii* and *S. bacillaris* have be found to have a respiratory quotient higher than *S. cerevisiae* (Quirós et al., 2014). Therefore, it seems that those are not respiratory yeasts that lower ethanol by respiration. The fact that simultaneous fermentations generally lower ethanol more that sequential inoculations (Fig. 3A) may indicate that the main effect of those yeast is the modulation of *S. cerevisiae* fermentative performance. Therefore, it was important to see the effect on *S. cerevisiae* transcriptome.

**Table 3.** Oenological parameters and main volatile compounds of experimental wines obtained from mixed starters cultures of selected non-*Saccharomyces* strains (Db2, Ha3, Ha9, St1, St8, St2, St5 and Zb1) sequentially inoculated with *S. cerevisiae* EC1118 (Sc). Pure culture of *S. cerevisiae* EC1118 (Control Sc) was used as control.

| Oenological characteristics | Db2+Sc | Ha3+Sc | Ha9+Sc | St1+Sc | St2+Sc | St5+Sc | St8+Sc | Zb1+Sc | Control (Sc) |
| --- | --- | --- | --- | --- | --- | --- | --- | --- | --- |
| Ethanol | 11.46±0.08<sup>a</sup> | 11.61±0.07<sup>a</sup> | 11.66±0.05<sup>a</sup> | 11.37±0.01<sup>ab</sup> | 11.70±0.13<sup>a</sup> | 11.54±0.09<sup>a</sup> | 11.34±0.13<sup>ab</sup> | 11.53±0.00<sup>a</sup> | 11.79±0.20<sup>a</sup> |
| Fructose | 2.00±0.14<sup>ab</sup> | 1.45±0.07<sup>a</sup> | 1.40±0.00<sup>a</sup> | 1.35±0.07<sup>a</sup> | 1.45±0.21<sup>a</sup> | 1.35±0.07<sup>a</sup> | 1.40±0.00<sup>a</sup> | 1.75±0.07<sup>a</sup> | 2.80±0.71<sup>b</sup> |
| Glucose | 2.10±0.00<sup>ad</sup> | 1.05±0.07<sup>bd</sup> | 1.15±0.07<sup>bc</sup> | 1.10±0.14<sup>bd</sup> | 1.05±0.35<sup>bd</sup> | 1.00±0.14<sup>bd</sup> | 0.95± 0.07<sup>bd</sup> | 1.80±0.28<sup>ac</sup> | 1.70±0.14<sup>cd</sup> |
| Total acidity | 5.70±0.06<sup>ab</sup> | 5.88±0.03<sup>ac</sup> | 5.98±0.03<sup>ac</sup> | 5.35±0.16<sup>bd</sup> | 5.73±0.12<sup>ab</sup> | 5.52±0.03<sup>ad</sup> | 5.78±0.25<sup>ab</sup> | 6.30±0.09<sup>c</sup> | 5.18±0.12<sup>d</sup> |
| Volatile acidity | 1.18±0.01<sup>a</sup> | 1.03±0.04<sup>ac</sup> | 0.69±0.01<sup>b</sup> | 0.88±0.06<sup>c</sup> | 0.97±0.04<sup>c</sup> | 1.17±0.01<sup>ad</sup> | 1.16±0.01<sup>ad</sup> | 0.91±0.10<sup>c</sup> | 0.99±0.05<sup>cd</sup> |
| Acetaldehyde | 46.26±3.92<sup>ab</sup> | 48.42±8.63<sup>ab</sup> | 33.11±3.61<sup>a</sup> | 27.75±0.28<sup>a</sup> | 39.46±1.31<sup>a</sup> | 38.32±10.33<sup>a</sup> | 42.96±6.36<sup>ab</sup> | 64.38±2.09<sup>b</sup> | 39.25±2.85<sup>a</sup> |
| Ethyl acetate | 28.14±6.98<sup>a</sup> | 13.45±0.33<sup>bc</sup> | 17.81±2.33<sup>c</sup> | 6.85±0.20<sup>b</sup> | 6.91±0.06<sup>b</sup> | 8.43±0.44<sup>bc</sup> | 8.88±1.17<sup>bc</sup> | 9.97±0.01<sup>bc</sup> | 10.09±1.29<sup>bc</sup> |
| *n*-Propanol | 28.54±0.78<sup>ab</sup> | 28.68±0.37<sup>ab</sup> | 31.76±2.16<sup>b</sup> | 22.73±3.34<sup>a</sup> | 22.33±3.34<sup>a</sup> | 24.74±0.73<sup>ab</sup> | 20.69±2.24<sup>a</sup> | 22.13±2.83<sup>a</sup> | 20.24±0.67<sup>a</sup> |
| Isobutanol | 29.70±0.51 | 26.24±1.06 | 30.91±8.37 | 33.14±0.07 | 33.07±8.64 | 23.46±0.34 | 24.38±0.32 | 32.80±4.00 | 32.15±2.08 |
| amyl alcohol | 40.78±2.70<sup>ac</sup> | 43.50±0.50<sup>a</sup> | 46.81±4.65<sup>a</sup> | 29.46±5.72<sup>bc</sup> | 27.06±0.50<sup>b</sup> | 33.13±0.53<sup>bd</sup> | 33.39±3.10<sup>bd</sup> | 42.27±2.70<sup>ad</sup> | 48.53±0.39<sup>a</sup> |
| Isoamyl alcohol | 102.66±0.14<sup>ab</sup> | 94.42±2.55<sup>ab</sup> | 104.50±1.07<sup>ab</sup> | 80.75±12.44<sup>a</sup> | 79.04±21.57<sup>a</sup> | 82.30±6.57<sup>a</sup> | 83.04±9.35<sup>a</sup> | 115.62±7.14<sup>ab</sup> | 127.23±2.59<sup>b</sup> |

Data are means ± standard deviation of two independent experiments. Different superscript letters in the same row correspond to statistically significant differences (Tukey's test, $p < 0.05$ ) between mixed and control fermentation. The oenological parameters are expressed as g/L, with exception of ethanol, expressed as % v/v; volatile compounds are expressed as mg/L

The impact of *S. bacillaris* in *S. cerevisiae* transcriptome is more conspicuous than the one of *Z. bailii* (Fig. 7). Interestingly, in the presence of both strains, *S. cerevisiae* increases the transcription of genes related to hexose metabolism (Table 4), up-regulating most of glycolytic genes. This behaviour may be seen being in contradiction with an ethanol decrease, but it may indicate that *S. cerevisiae* is struggling to achieve all the energy required and it is gearing up this pathway. A similar behaviour in glycolysis transcription has been seen in mixed fermentations with *M. pulcherrima* in aerobic conditions (Mencher et al., 2021), although this yeast represses also aerobic respiration, unlike the two species tested in synthetic grape juice fermentation. Stimulation of nutrient consumption seems to be a common trend in co-cultivation with many non-conventional yeast (Curiel et al., 2017). However, there are some molecular signatures that may indicate specific ways of regulation of carbon metabolism in our selected strains. In the presence of *Z. bailli, S. cerevisiae* increase significantly the GO categories of trehalose and glycogen synthesis, inducing the main genes involved in trehalose (trehalose synthase genes *TPS1, TPS2, TSL1*, Supplementary Table S2) and glycogen (glycogen synthase genes *GSY1* and *GSY2*). Genes involved in glycogen degradation like glycogen debranching *GDB1* and glycogen phosphorylase *GPH1* are also induced, suggesting a potential futile cycle. If *S. cerevisiae* diverts part of glucose to the accumulation of reserve and stress tolerance polysaccharides, that may reduce the amount of glucose to be fermented into ethanol. In fact overexpression of trehalose synthesis in wine yeasts leads to reduction of ethanol (Rossouw et al., 2013). *S. bacillaris* increases in a specific way the genes of the GO “gluconeogenesis” (Table 4). Those are basically glycolytic genes, but gluconeogenic specific genes like pyruvate carboxylase *PYC1*, phosphoenolpyruvate carboxykinase *PCK1* and fructose-1,6-bisphosphatase *FBP1* are up-regulated, and that may be causing two futile cycles that may be consuming energy with no gain, reducing the ethanol yield per glucose molecule. A common glycolytic enzyme induced by the two yeasts is the glyceraldehyde 3-phosphate dehydrogenase coded by *TDH1-3* genes. This enzyme has been defined as a cell wall component (ML et al., 2001) and some of it fragments have been described as antimicrobial peptides against a variety of yeasts (Branco et al., 2014), so it may be behind death of the non-*Saccharomyces* yeast, particularly *S. bacillaris* by cell-to-cell interactions (Englezos et al., 2019b).

**Table 4.** Main Biological Process GO categories over-represented in mixed fermentations. See Supplementary Table S3 for a full analysis.

| Fermentation | GO categories | Fold enrichment |
| --- | --- | --- |
| **Zb+Sc vs Sc, up-regulated** | 'de novo' NAD biosynthetic process | 13,91 |
|  | glycolytic process | 10.43 |
|  | trehalose metabolic process | 9.48 |
|  | glycogen metabolic process | 5.62 |
| **Zb+Sc vs Sc, down-regulated** | None |  |
| **Sb+Sc vs Sc, up-regulated** | gluconeogenesis (GO:0006094) | 7.62 |
|  | rRNA transport | 6.38 |
|  | cytoplasmic translation | 5.38 |
| **Sb+Sc vs Sc, down-regulated** | cytoplasm to vacuole transport by the Cvt pathway | 4.11 |
|  | protein targeting to vacuole | 2.92 |
| **Zb+Sc vs Sb+Sc, up-regulated** | late nucleophagy | 7.01 |
|  | homoserine metabolic process | 6.95 |
|  | iron ion transmembrane transport | 6.95 |
| **Zb+Sc vs Sb+Sc, down-regulated** | rRNA export from nucleus | 9.62 |
|  | cytoplasmic translation | 8.61 |

The transcriptomic analysis reveals some specific interactions. For instance, the presence of *S. bacillaris* repress genes involved in iron assimilation. Interestingly genes of siderophores (*ARN1, ARN2, SIT1*) and particularly two genes of cell wall mannoproteins involved in the retention of siderophore-iron in the cell wall, *FIT2* and *FIT3* (Fig. 7C) are among them. In synthetic grape juice there are no siderophores, and it is known that *S. cerevisiae* do not produce them (Martínez-Pastor and Puig, 2020). However, it has been reported the acquisition by some *Starmerella* yeasts of a bacterial operon to synthetize the siderophore enterobactin by horizontal gene transfer (Kominek et al., 2019). The presence of an excess of siderophore may be detrimental, as an excess of iron could trigger oxidative stress, so *S. cerevisiae* prevents its import. The upregulation of *S. cerevisae*’s protein synthesis and down regulation of vesicular transport by the presence of *S. bacillaris* are indicating a profound rearrangement of biosynthetic processes that would require a careful and complex analysis.

## Acknowledgements

This work was funded by a grant from the Spanish Ministry of Science (AGL2017-83254-R) to EM and AA. This work was supported by the projects PSR Regione Basilicata 2014-2020, sottomisura 16.1 GO Vites&Vino PROduttività e Sostenibilità in vITivinicoltura (PROSIT), N. 54250365779 and sottomisura 16.2 IN.VINI.VE.RI.TA.S (Innovare la viti-VINIcoltura lucana: VErso la RIgenerazione varieTAle, la Selezione di vitigni locali e proprietà antiossidanti dei vini), N. 976 to AC. Furthermore, the JRU MIRRI-IT (http://www.mirri-it.it/) is greatly acknowledged for scientific support.

## References

Benito, Á., Calderón, F., Benito, S., 2019. The influence of non-saccharomyces species on wine fermentation quality parameters. Fermentation. https://doi.org/10.3390/fermentation5030054

Branco, P., Francisco, D., Chambon, C., Hébraud, M., Arneborg, N., Almeida, M.G., Caldeira, J., Albergaria, H., 2014. Identification of novel GAPDH-derived antimicrobial peptides secreted by Saccharomyces cerevisiae and involved in wine microbial interactions. Appl. Microbiol. Biotechnol. 98, 843–853. https://doi.org/10.1007/s00253-013-5411-y

Capece, A., Romano, P., 2019. Yeasts and Their Metabolic Impact on Wine Flavour BT-Yeasts in the Production of Wine, in: Romano, P., Ciani, M., Fleet, G.H. (Eds.),. Springer New York, New York, NY, pp. 43–80. https://doi.org/10.1007/978-1-4939-9782-4_2

Capece, A., Siesto, G., Poeta, C., Pietrafesa, R., Romano, P., 2013. Indigenous yeast population from Georgian aged wines produced by traditional “Kakhetian” method. Food Microbiol. 36, 447–455. https://doi.org/https://doi.org/10.1016/j.fm.2013.07.008

Ciani, M., Capece, A., Comitini, F., Canonico, L., Siesto, G., Romano, P., 2016. Yeast Interactions in Inoculated Wine Fermentation. Front. Microbiol. 7, 555. https://doi.org/10.3389/fmicb.2016.00555

Contreras, A., Hidalgo, C., Schmidt, S., Henschke, P.A., Curtin, C., Varela, C., 2015. The application of non-Saccharomyces yeast in fermentations with limited aeration as a strategy for the production of wine with reduced alcohol content. Int. J. Food Microbiol. 205, 7–15. https://doi.org/10.1016/j.ijfoodmicro.2015.03.027

Curiel, J.A., Morales, P., Gonzalez, R., Tronchoni, J., 2017. Different Non-Saccharomyces Yeast Species Stimulate Nutrient Consumption in S. cerevisiae Mixed Cultures. Front. Microbiol..

De Orduna, R.M., 2010. Climate change associated effects on grape and wine quality and production. Food Res. Int. 43, 1844–1855.

Englezos, V., Pollon, M., Rantsiou, K., Ortiz-Julien, A., Botto, R., Río Segade, S., Giacosa, S., Rolle, L., Cocolin, L., 2019a. Saccharomyces cerevisiae-Starmerella bacillaris strains interaction modulates chemical and volatile profile in red wine mixed fermentations. Food Res. Int. 122, 392–401. https://doi.org/10.1016/j.foodres.2019.03.072

Englezos, V., Rantsiou, K., Cravero, F., Torchio, F., Ortiz-Julien, A., Gerbi, V., Rolle, L., Cocolin, L., 2016. Starmerella bacillaris and Saccharomyces cerevisiae mixed fermentations to reduce ethanol content in wine. Appl. Microbiol. Biotechnol. 100, 5515–5526. https://doi.org/10.1007/s00253-016-7413-z

Englezos, V., Rantsiou, K., Giacosa, S., Río Segade, S., Rolle, L., Cocolin, L., 2019b. Cell-to-cell contact mechanism modulates Starmerella bacillaris death in mixed culture fermentations with Saccharomyces cerevisiae. Int. J. Food Microbiol. 289, 106–114. https://doi.org/10.1016/j.ijfoodmicro.2018.09.009

Escott, C., Del Fresno, J.M., Loira, I., Morata, A., Suárez-Lepe, J.A., 2018. Zygosaccharomyces rouxii: Control Strategies and Applications in Food and Winemaking. Fermentation 4, 69.

Fleet, G.H., 1993. Wine microbiology and biotechnology. Harwood Academic Publishers, Chur; Philadelphia, Pa.

Gobbi, M., De Vero, L., Solieri, L., Comitini, F., Oro, L., Giudici, P., Ciani, M., 2014. Fermentative aptitude of non-Saccharomyces wine yeast for reduction in the ethanol content in wine. Eur. Food Res. Technol. 239, 41–48. https://doi.org/10.1007/s00217-014-2187-y

Gonzalez, R., Quirós, M., Morales, P., 2013. Yeast respiration of sugars by non-Saccharomyces yeast species: A promising and barely explored approach to lowering alcohol content of wines. Trends Food Sci. Technol. https://doi.org/10.1016/j.tifs.2012.06.015

Granchi, L., Bosco, M., Messini, A., Vincenzini, M., 1999. Rapid detection and quantification of yeast species during spontaneous wine fermentation by PCR-RFLP analysis of the rDNA ITS region. J. Appl. Microbiol. 87, 949–956. https://doi.org/10.1046/j.1365-2672.1999.00600.x

Hranilovic, A., Li, S., Boss, P.K., Bindon, K., Ristic, R., Grbin, P.R., Van der Westhuizen, T., Jiranek, V., 2018. Chemical and sensory profiling of Shiraz wines co-fermented with commercial non-Saccharomyces inocula. Aust. J. Grape Wine Res. 24, 166–180. https://doi.org/https://doi.org/10.1111/ajgw.12320

Jolly, N.P., Varela, C., Pretorius, I.S., 2014. Not your ordinary yeast: non-Saccharomyces yeasts in wine production uncovered. FEMS Yeast Res 14, 215–237. https://doi.org/10.1111/1567-1364.12111

Kominek, J., Doering, D.T., Opulente, D.A., Shen, X.-X., Zhou, X., DeVirgilio, J., Hulfachor, A.B., Groenewald, M., Mcgee, M.A., Karlen, S.D., Kurtzman, C.P., Rokas, A., Hittinger, C.T., 2019. Eukaryotic Acquisition of a Bacterial Operon. Cell 176, 1356–1366.e10. https://doi.org/10.1016/j.cell.2019.01.034

Lemos Junior, W.J.F., da Silva Duarte, V., Treu, L., Campanaro, S., Nadai, C., Giacomini, A., Corich, V., 2018. Whole genome comparison of two Starmerella bacillaris strains with other wine yeasts uncovers genes involved in modulating important winemaking traits. FEMS Yeast Res. 18, foy069.

Lemos Junior, W.J.F., de Oliveira, V.S., Guerra, A.F., Giacomini, A., Corich, V., 2021. From the vineyard to the cellar: new insights of Starmerella bacillaris (synonym Candida zemplinina) technological properties and genomic perspective. Appl. Microbiol. Biotechnol. 105, 493–501. https://doi.org/10.1007/s00253-020-11041-9

Love, M.I., Huber, W., Anders, S., 2014. Moderated estimation of fold change and dispersion for RNA-seq data with DESeq2. Genome Biol. 15, 550. https://doi.org/10.1186/s13059-014-0550-8

Manzanares, P., Ramón, D., Querol, A., 1999. Screening of non-Saccharomyces wine yeasts for the production of β-D-xylosidase activity. Int. J. Food Microbiol. 46, 105–112. https://doi.org/https://doi.org/10.1016/S0168-1605(98)00186-X

Manzanares, P., Rojas, V., Genovés, S., Vallés, S., 2000. A preliminary search for anthocyanin-β-D-glucosidase activity in non-Saccharomyces wine yeasts. Int. J. Food Sci. Technol. 35, 95–103. https://doi.org/https://doi.org/10.1046/j.1365-2621.2000.00364.x

Martin, V., Valera, M.J., Medina, K., Boido, E., Carrau, F., 2018. Oenological Impact of the Hanseniaspora/Kloeckera Yeast Genus on Wines—A Review. Fermentation 4, 76.

Martínez-Pastor, M.T., Puig, S., 2020. Adaptation to iron deficiency in human pathogenic fungi. Biochim. Biophys. Acta - Mol. Cell Res. 1867, 118797. https://doi.org/https://doi.org/10.1016/j.bbamcr.2020.118797

Masneuf-Pomarede, I., Bely, M., Marullo, P., Albertin, W., 2016. The Genetics of Non-conventional Wine Yeasts: Current Knowledge and Future Challenges. Front. Microbiol. 6, 1563. https://doi.org/10.3389/fmicb.2015.01563

Medina, K., Boido, E., Dellacassa, E., Carrau, F., 2012. Growth of non-Saccharomyces yeasts affects nutrient availability for Saccharomyces cerevisiae during wine fermentation. Int. J. Food Microbiol. 157, 245–250. https://doi.org/10.1016/j.ijfoodmicro.2012.05.012

Medina, K., Boido, E., Farina, L., Gioia, O., Gomez, M.E., Barquet, M., Gaggero, C., Dellacassa, E., Carrau, F., 2013. Increased flavour diversity of Chardonnay wines by spontaneous fermentation and co-fermentation with Hanseniaspora vineae. Food Chem 141, 2513–2521. https://doi.org/10.1016/j.foodchem.2013.04.056

Mencher, A., Morales, P., Curiel, J.A., Gonzalez, R., Tronchoni, J., 2021. Metschnikowia pulcherrima represses aerobic respiration in Saccharomyces cerevisiae suggesting a direct response to co-cultivation. Food Microbiol. 94, 103670. https://doi.org/10.1016/j.fm.2020.103670

Mi, H., Muruganujan, A., Ebert, D., Huang, X., Thomas, P.D., 2019. PANTHER version 14: more genomes, a new PANTHER GO-slim and improvements in enrichment analysis tools. Nucleic Acids Res. 47, D419–D426. https://doi.org/10.1093/nar/gky1038

Milanovic, V., Ciani, M., Oro, L., Comitini, F., 2012. Starmerella bombicola influences the metabolism of Saccharomyces cerevisiae at pyruvate decarboxylase and alcohol dehydrogenase level during mixed wine fermentation. Microb. Cell Fact. 11, 18. https://doi.org/10.1186/1475-2859-11-18

Ml, D., Je, O., Azori, N.I., Renau-Piqueras, J., Ml, G., Gozalbo, D., 2001. The glyceraldehyde-3-phosphate dehydrogenase polypeptides encoded by the Saccharomyces cerevisiae TDH1, TDH2 and TDH3 genes are also cell wall proteins. Microbiology 147, 411–417.

Nissen, P., Nielsen, D., Arneborg, N., 2003. Viable Saccharomyces cerevisiae cells at high concentrations cause early growth arrest of non-Saccharomyces yeasts in mixed cultures by a cell-cell contact-mediated mechanism. Yeast 20, 331–341. https://doi.org/10.1002/yea.965

Orlova, M., Barrett, L., Kuchin, S., 2008. Detection of endogenous Snf1 and its activation state: application to Saccharomyces and Candida species. Yeast 25, 745–754. https://doi.org/10.1002/yea.1628

Padilla, B., Gil, J. V, Manzanares, P., 2016. Past and Future of Non-Saccharomyces Yeasts: From Spoilage Microorganisms to Biotechnological Tools for Improving Wine Aroma Complexity. Front. Microbiol. 7. https://doi.org/10.3389/fmicb.2016.00411

Patro, R., Duggal, G., Love, M.I., Irizarry, R.A., Kingsford, C., 2017. Salmon provides fast and bias-aware quantification of transcript expression. Nat. Methods 14, 417–419. https://doi.org/10.1038/nmeth.4197

Pérez-Torrado, R., Gamero, E., Gómez-Pastor, R., Garre, E., Aranda, A., Matallana, E., 2015. Yeast biomass, an optimised product with myriad applications in the food industry. Trends Food Sci. Technol. 46, 167–175. https://doi.org/http://dx.doi.org/10.1016/j.tifs.2015.10.008

Petruzzi, L., Capozzi, V., Berbegal, C., Corbo, M.R., Bevilacqua, A., Spano, G., Sinigaglia, M., 2017. Microbial Resources and Enological Significance: Opportunities and Benefits. Front. Microbiol. 8, 995. https://doi.org/10.3389/fmicb.2017.00995

Quirós, M., Rojas, V., Gonzalez, R., Morales, P., 2014. Selection of non-Saccharomyces yeast strains for reducing alcohol levels in wine by sugar respiration. Int. J. Food Microbiol. https://doi.org/10.1016/j.ijfoodmicro.2014.04.024

Renault, P.E., Albertin, W., Bely, M., 2013. An innovative tool reveals interaction mechanisms among yeast populations under oenological conditions. Appl. Microbiol. Biotechnol. 97, 4105–4119. https://doi.org/10.1007/s00253-012-4660-5

Ribéreau-Gayon, P., Dubourdieu, D., Donèche, B., 2006. Handbook of enology, 2nd ed. John Wiley, Chichester, West Sussex, England; Hoboken, NJ.

Rodicio, R., Heinisch, J.J., 2009. Sugar Metabolism by Saccharomyces and non-Saccharomyces Yeasts BT - Biology of Microorganisms on Grapes, in Must and in Wine, in: König, H., Unden, G., Fröhlich, J. (Eds.),. Springer Berlin Heidelberg, Berlin, Heidelberg, pp. 113–134. https://doi.org/10.1007/978-3-540-85463-0_6

Rossouw, D., Heyns, E.H., Setati, M.E., Bosch, S., Bauer, F.F., 2013. Adjustment of Trehalose Metabolism in Wine Saccharomyces cerevisiae Strains To Modify Ethanol Yields. Appl. Environ. Microbiol. 79, 5197–5207. https://doi.org/10.1128/aem.00964-13

Roudil, L., Russo, P., Berbegal, C., Albertin, W., Spano, G., Capozzi, V., 2019. Non-Saccharomyces Commercial Starter Cultures: Scientific Trends, Recent Patents and Innovation in the Wine Sector. Recent Pat. Food. Nutr. Agric. https://doi.org/10.2174/2212798410666190131103713

Torrellas, M., Rozès, N., Aranda, A., Matallana, E., 2020. Basal catalase activity and high glutathione levels influence the performance of non-Saccharomyces active dry wine yeasts. Food Microbiol. https://doi.org/10.1016/j.fm.2020.103589

Vallejo, B., Matallana, E., Aranda, A., 2020. Saccharomyces cerevisiae nutrient signaling pathways show an unexpected early activation pattern during winemaking. Microb. Cell Fact. https://doi.org/10.1186/s12934-020-01381-6

Wang, C., Esteve-Zarzoso, B., Mas, A., 2014. Monitoring of Saccharomyces cerevisiae, Hanseniaspora uvarum, and Starmerella bacillaris (synonym Candida zemplinina) populations during alcoholic fermentation by fluorescence in situ hybridization. Int. J. Food Microbiol. 191, 1–9. https://doi.org/https://doi.org/10.1016/j.ijfoodmicro.2014.08.014

Wang, C., Mas, A., Esteve-Zarzoso, B., 2016. The Interaction between Saccharomyces cerevisiae and Non-Saccharomyces Yeast during Alcoholic Fermentation Is Species and Strain Specific. Front. Microbiol..

Zhu, X., Navarro, Y., Mas, A., Torija, M.-J., Beltran, G., 2020. A Rapid Method for Selecting Non-Saccharomyces Strains with a Low Ethanol Yield. Microorg.. https://doi.org/10.3390/microorganisms8050658

