## Supplementary Table S1 for "Impact of *Starmerella bacillaris* and *Zygosaccharomyces bailii* on ethanol reduction and *Saccharomyces cerevisiae* metabolism during mixed wine fermentations"

**Supplementary Table S1.** Concentration of major chemical compounds of experimental wines obtained from mixed starters cultures of selected non-*Saccharomyces* strains (St8 and Zb1) simultaneous inoculated with *S. cerevisiae* EC1118 (Sc). Pure culture of *S. cerevisiae* EC1118 (Control Sc) was used as a control.

| **Oenological characteristics** | **St8+Sc** | **Zb1+Sc** | **Control (Sc)** |
|---|---|---|---|
| Ethanol % (v/v) | 13.44±0.0[a] | 13.07±0.1[a] | 13.92±0.3[b] |
| Fructose (g/L) | 1.43±0.3[a] | 1.90±1.0[a] | 4.20±1.0[b] |
| Glucose (g/L) | 1.0±0.1 | 1.33±0.3 | 1.10±0.3 |
| Total acidity (g/L) | 6.30±0.1 | 6.60±0.3 | 6.26±0.2 |
| Volatile acidity (g/L) | 0.59±0.1 | 0.82±0.4 | >1.0 |
| pH | 3.50±0.0 | 3.54±0.0 | 3.51±0.0 |
| Malic acid (g/L) | 1.60±0.1 | 1.60±0.1 | 1.73±0.1 |
| Lactic acid (g/L) | 0.00±0.0 | 0.00±0.0 | 0.00±0.0 |
| **Main volatile compounds** | | | |
| Acetaldehyde (mg/L) | 45.34±3.26[a] | 42.38±3.58[a] | 73.37±6.47[b] |
| Ethyl acetate (mg/L) | 23.32±1.98 | 23.74±3.48 | 20.65±1.85 |
| *n*-Propanol (mg/L) | 29.94±0.75[a] | 34.18±2.38[ab] | 39.76±5.72[b] |
| Isobutanol (mg/L) | 21.58±1.82[a] | 20.81±1.67[a] | 15.92±1.79[b] |
| *n*-Butanol (mg/L) | 12.00±1.41[a] | 16.70±4.45[a] | 24.93±3.18[b] |
| Acetoin (mg/L) | 3.80±0.62[a] | 4.57±0.50[a] | 8.90±0.56[b] |
| D-amyl alcohol (mg/L) | 165.91±3.49[a] | 22.43±0.65[b] | 28.19±4.84[b] |
| Isoamyl alcohol (mg/L) | 51.40±9.00[a] | 37.20±4.71[ab] | 35.60±1.68[b] |

Note: Data are means ± standard deviation of three independent experiments. Different superscript letters in the same row correspond to statistically significant differences (Tukey's test, $p < 0.05$).
