## Supplementary Figure S1 for "Impact of *Starmerella bacillaris* and *Zygosaccharomyces bailii* on ethanol reduction and *Saccharomyces cerevisiae* metabolism during mixed wine fermentations"

A)

B)

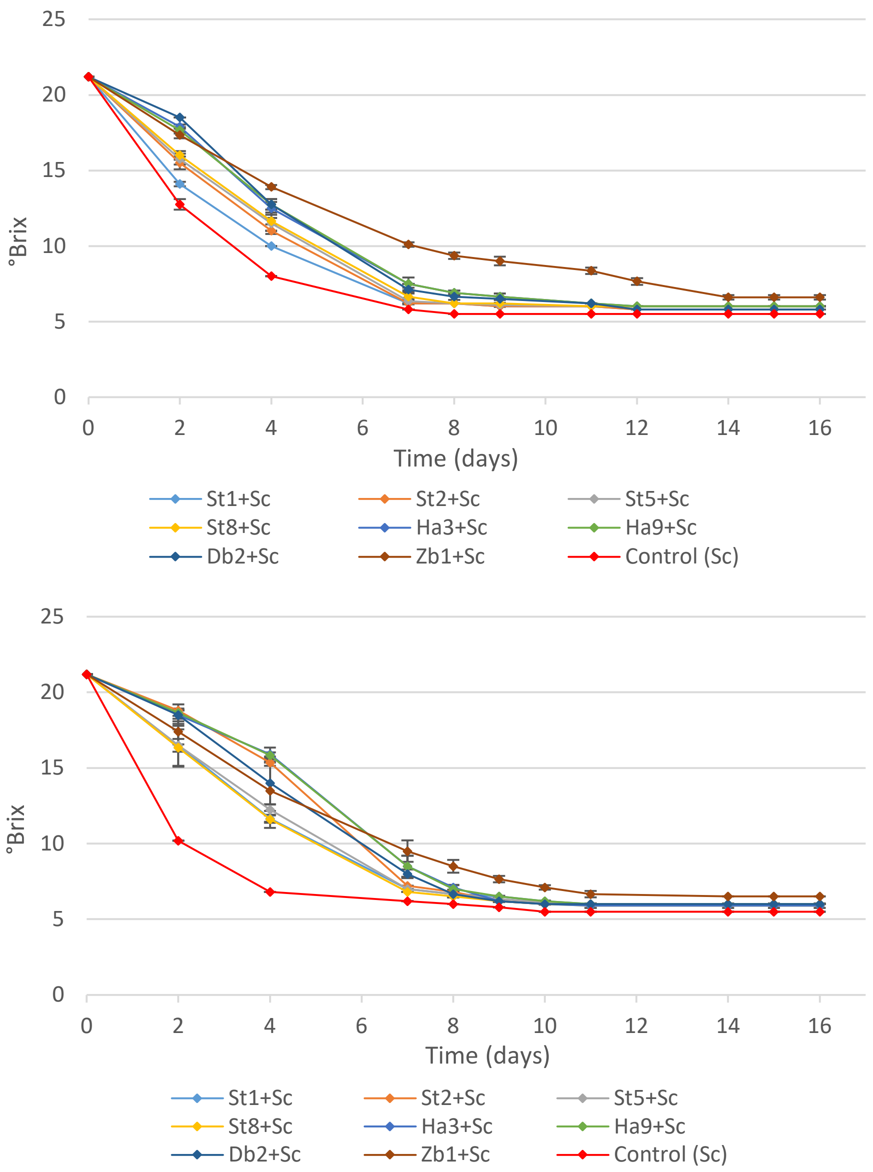


**Supplementary Figure S1**. Sugar consumption of mixed starters cultures of *D. polymorphus* (Db2), *H. uvarum* (Ha3 and Ha9), *St. bacillaris* (St1, St8, St2 and St5) and *Z. bailii* (Zb1) strains simultaneously A) or sequentially B) inoculated with *S. cerevisiae* EC1118 (Sc). Pure culture of *S. cerevisiae* EC1118 (Control Sc) was used as a control. Data are means ± standard deviation of two independent experiments
