## Supplementary Figure S2 for "Impact of *Starmerella bacillaris* and *Zygosaccharomyces bailii* on ethanol reduction and *Saccharomyces cerevisiae* metabolism during mixed wine fermentations"

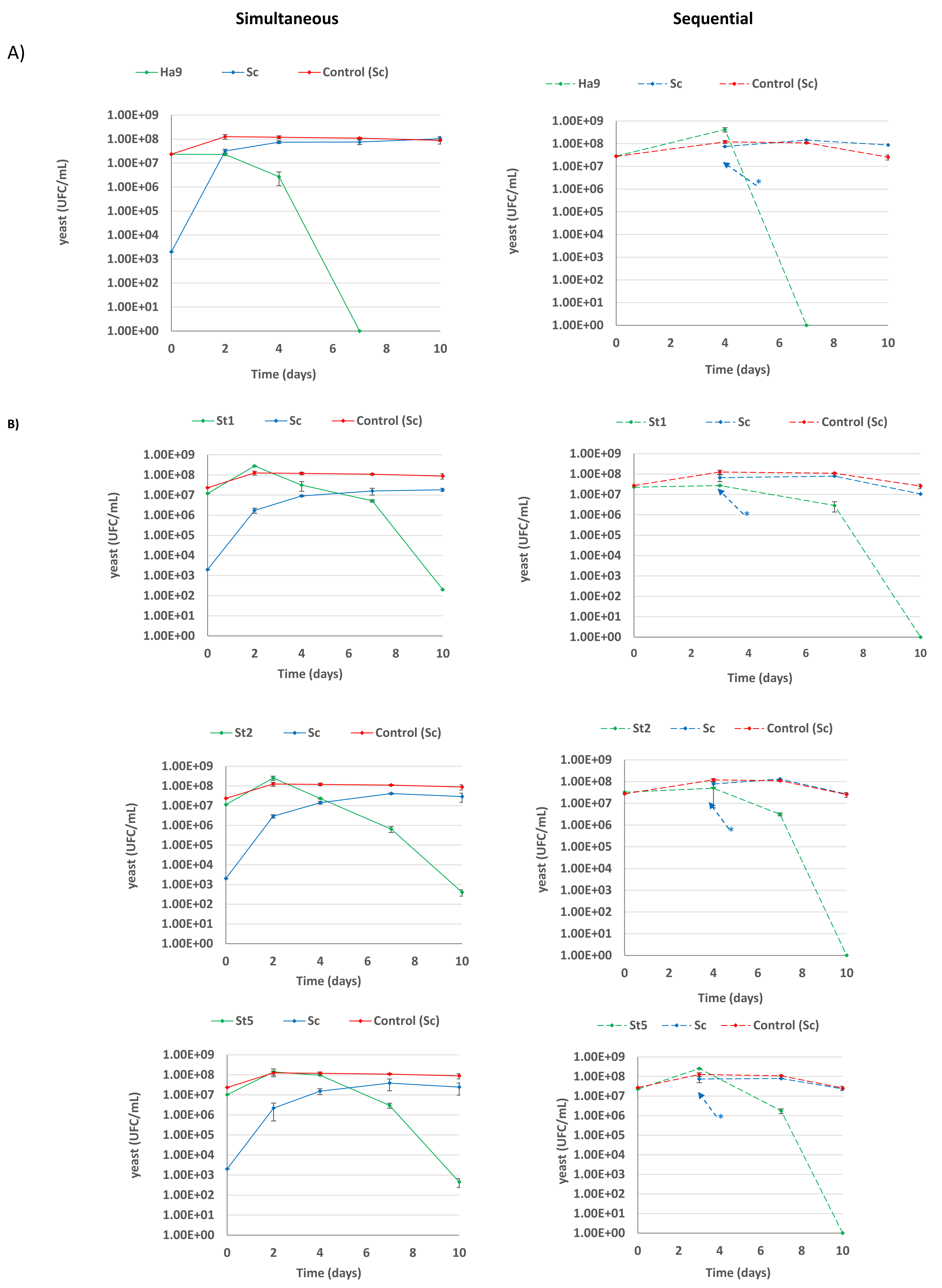


**Supplementary Fig. S2**. Evolution of yeast populations in mixed fermentations inoculated with *S. cerevisiae* (Sc) and *H. uvarum* (Ha9, A)) and *S. bacillaris* (St1, St2, St5, B)) in simultaneous and sequential modalities. Values are mean of two independent duplicates. Pure culture of *S. cerevisiae* EC1118 (Control Sc) was used as a control. * indicate the inoculation time of *S. cerevisiae.*
